# Phase-Separating Peptide Coacervates with Programmable Material Properties for Universal Intracellular Delivery of Macromolecules

**DOI:** 10.1101/2024.06.20.599859

**Authors:** Yue Sun, Xi Wu, Jianguo Li, Milad Radiom, Raffaele Mezzenga, Chandra Shekhar Verma, Jing Yu, Ali Miserez

## Abstract

Phase-separating peptides (PSPs) self-assembling into coacervate microdroplets (CMs) are a promising new class of intracellular delivery vehicles that can release macromolecular modalities deployed in a wide range of therapeutic treatments. However, the molecular grammar governing intracellular uptake and release kinetics of CMs remains elusive. Here, we systematically manipulated the sequence of PSPs to unravel the relationships between their molecular structure, the physical properties of the resulting CMs, and their delivery efficacy. We show that a few amino acid alterations are sufficient to modulate the viscoelastic properties of CMs towards either a gel-like or a liquid-like state as well as their binding interaction with cellular membranes, collectively enabling to tune the kinetics of intracellular cargo release. Our findings provide molecular guidelines to precisely program the material properties of PSP CMs and achieve tunable cellular uptake and release kinetics depending on the cargo modality, with broad implications for therapeutic applications.

## Introduction

Efficient and safe intracellular delivery of macromolecules stands as a pivotal requirement for promising therapeutic avenues, spanning protein-, gene-, and immuno-therapies.(*1–6*) However, current methodologies, including viral vectors, electroporation, and nanocarriers, grapple with persistent concerns such as long-term safety, cytotoxicity, low cargo encapsulation efficiency, and manufacturing complexity.(*7–10*) Recently, we developed coacervate microdroplets (CMs) assembled from phase-separating peptides (PSPs) by pH-induced liquid-liquid phase separation (LLPS) as a simple, safe, and versatile approach capable of delivering a broad spectrum of macromolecular therapeutics into cells.(*11, 12*) Our PSPs, inspired after histidine-rich beak proteins (HBPs)(*13, 14*) and abbreviated HB*pep-*SP, are characterized by a modular design comprised of the consensus pentapeptide GHGXY, where X is a variable residue(*15, 16*). To confer redox-triggered cargo release capability upon HB*pep-*SP CMs, we introduced a single lysine (Lys) residue conjugated with a moiety reducible by endogenous glutathione (GSH) within the cytosol (Fig. S1). The resulting CMs can recruit various macromolecular therapeutics during LLPS and release them into cells, activated by GSH-induced disassembly of CMs.(*12*) Here, we demonstrate that the modular sequence of HB*pep*-SP enables the systematic exploration of the X position, allowing for the incorporation of diverse amino acids (Fig. 1). According to molecular dynamics (MD) simulations, this positional coding alters peptide-peptide interactions and molecular hydration, thereby enabling to modulate the viscoelastic properties of CMs from soft gels to viscous liquids. Furthermore, these mutations also control the binding interactions of CMs to the cell membrane, and collectively these properties regulate cargo release efficacy. We then show optimal X residue combinations that maximize transfection efficiency for various payloads, encompassing proteins, peptides, plasmid DNA (pDNA), messenger RNA (mRNA), small interfering RNA (siRNA), and CRISPR/Cas9 gene editing machinery. Moreover, we show that this excellent delivery efficiency can be extended towards hard-to-transfect cells such as primary and immune cells.

**Fig. 1.**
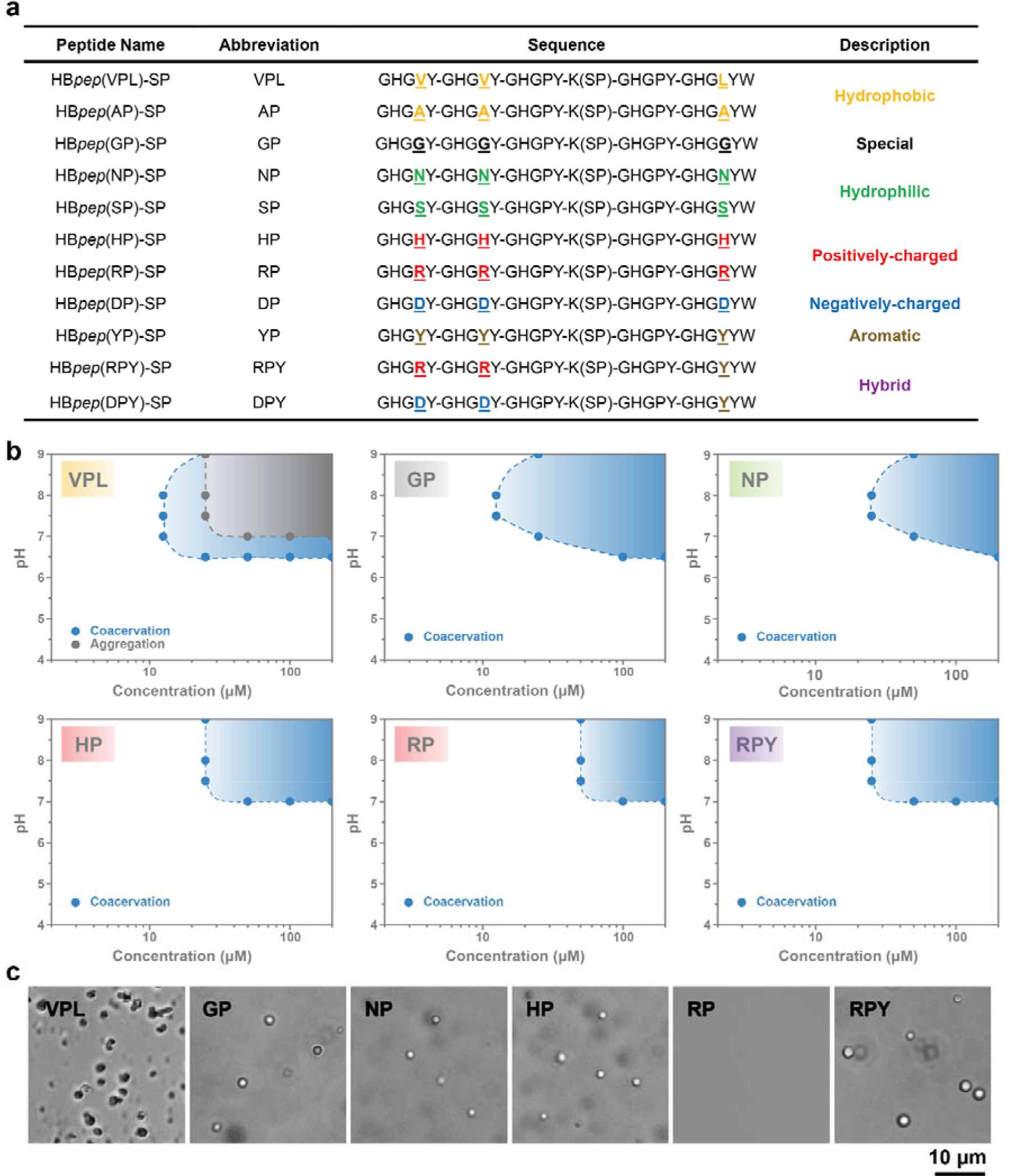
LLPS of HB*pep*-SP variants. (a) Peptide sequence of HB*pep*-SP variants. K(SP) represents the lysine (Lys, K) residue conjugated with the moiety cleaved by GSH as shown in Fig. S1. The abbreviations used in the text refer to the X1, X2 and X5 positions in the pentapeptide repeat GHGXY. When X1, X2 and X5 are the same residue, a two-letter abbreviation is used, with the first letter corresponding to that residue and the second letter to proline (Pro, P) since the X3 and X4 positions are P in all cases. When X5 position is different from X1 and X2, a three-letter abbreviation is used (first letter for position X1 and X2 and third letter for position X5). **(b)** Phase diagram of representative variants at the ionic strength (IS) of 100 mM, with the region of coacervation or aggregation shadowed in blue and grey, respectively. **(c)** Representative optical micrographs of HB*pep*-SP variants (50 μM, pH = 7.0, IS = 100 mM).

Overall, our findings firmly establish PSPs coacervate engineering as an intracellular delivery toolbox for diverse therapeutics, offering programmable efficiency tailored to cargo modality and cell type.

## Results

### LLPS of the HB*pep*-SP variants

HB*pep*-SPs consist of five repeats of the motif GHGXY, where X is a variable residue. A single Lys is placed after the third repeat and conjugated with a sidechain containing a disulfide and a self-immolative moieties, while tryptophan (W) is added at the C-terminus (Fig. 1a). Since valine (V), proline (P) and leucine (L) are found at the X positions in native HBPs, we previously designed HB*pep*(VPL)-SP (abbreviated VPL).(*12, 17*) However, the hydrophobic nature of V and L residues limited the phase behavior of VPL, with CMs forming only near pH 6.5 (Figs. 1b-c), posing limitations for therapeutics that may not be soluble at this pH. By substituting the three X positions with less hydrophobic glycine (G) and alanine (A) residues, the phase behavior could be altered: HB*pep*(GP)-SP and HB*pep*(AP)-SP (GP and AP) formed CMs across a broader pH range up to pH 9.0 (Figs. 1a-c and S2). Further substitutions with more hydrophilic residues asparagine (N) and serine (S) narrowed the area of two-phase regions (NP and SP, Figs. 1a-c and S2), underscoring the role of hydrophobic interactions during LLPS of PSPs.(*18–20*) We then explored variants incorporating the charged residues histidine (H), arginine (R) and aspartic acid (D) at the X positions. HB*pep*(HP)-SP (HP) and HB*pep*(RP)-SP (RP) only phase-separated above pH 7.0 owing to electrostatic repulsion from their positively-charged residues. In contrast, HB*pep*(DP)-SP (DP) phase-separated in the lower pH range 5.0 - 6.5 due to its lower isoelectric point. Owing to the histidine pKa of 6.5, HP is neutral above pH 7, enabling phase-separation to occur at lower peptide concentrations, whereas phase-separation at pH 7 for RP and DP requires a higher peptide concentration (Figs. 1b and S2). This effect could be counteracted by mutating the X positions with the aromatic tyrosine (Y), which enhanced hydrophobic and π-π stacking interactions.(*21*) Indeed, this YP peptide formed aggregates across a wide pH range from 6.5 to 9.0, indicative of enhanced inter-peptide interactions (Fig. S2). Consequently, two hybrid variants, RPY and DPY, were designed by replacing R and D at the X5 position with Y. RPY and DPY phase-separated over a broader range of conditions compared to RP and DP variants (Figs. 1b and S2), emphasizing the ability to precisely control LLPS conditions by simple amino acid mutations.

### Materials and biophysical properties of HB*pep*-SP CM variants

Having established the LLPS tunability of the HB*pep*-SP variants, we delved deeper into the material and biophysical properties of CMs. Utilizing the surface force apparatus (SFA) technique,(*22, 23*) we first assessed their viscoelastic characteristics. The SFA employs two mica surfaces affixed on perpendicularly oriented cylindrical discs, between which the samples are injected. The surfaces coated with the CMs were approached with sub-nm distance resolution and then retracted, while the normal force between the cylinders was monitored with a few μN force sensitivity. During separation, CMs formed by NP and GP (Fig. 2a) exhibited mechanical instability –indicated by a sudden “jump-out” on the normalized force-distance (*F-D*) curves– suggesting gel-like properties.(*24*) Conversely, VPL, HP, and RPY CMs exhibited *F*-*D* curves with continuous retraction forces, indicative of viscous liquid bridges between the surfaces.(*25*) Further evidence of the gel versus fluidic behavior was evident from the hysteresis during approach/separation cycles, which were markedly present for HP and RPY CMs but minimal for GP and NP.

**Fig. 2.**
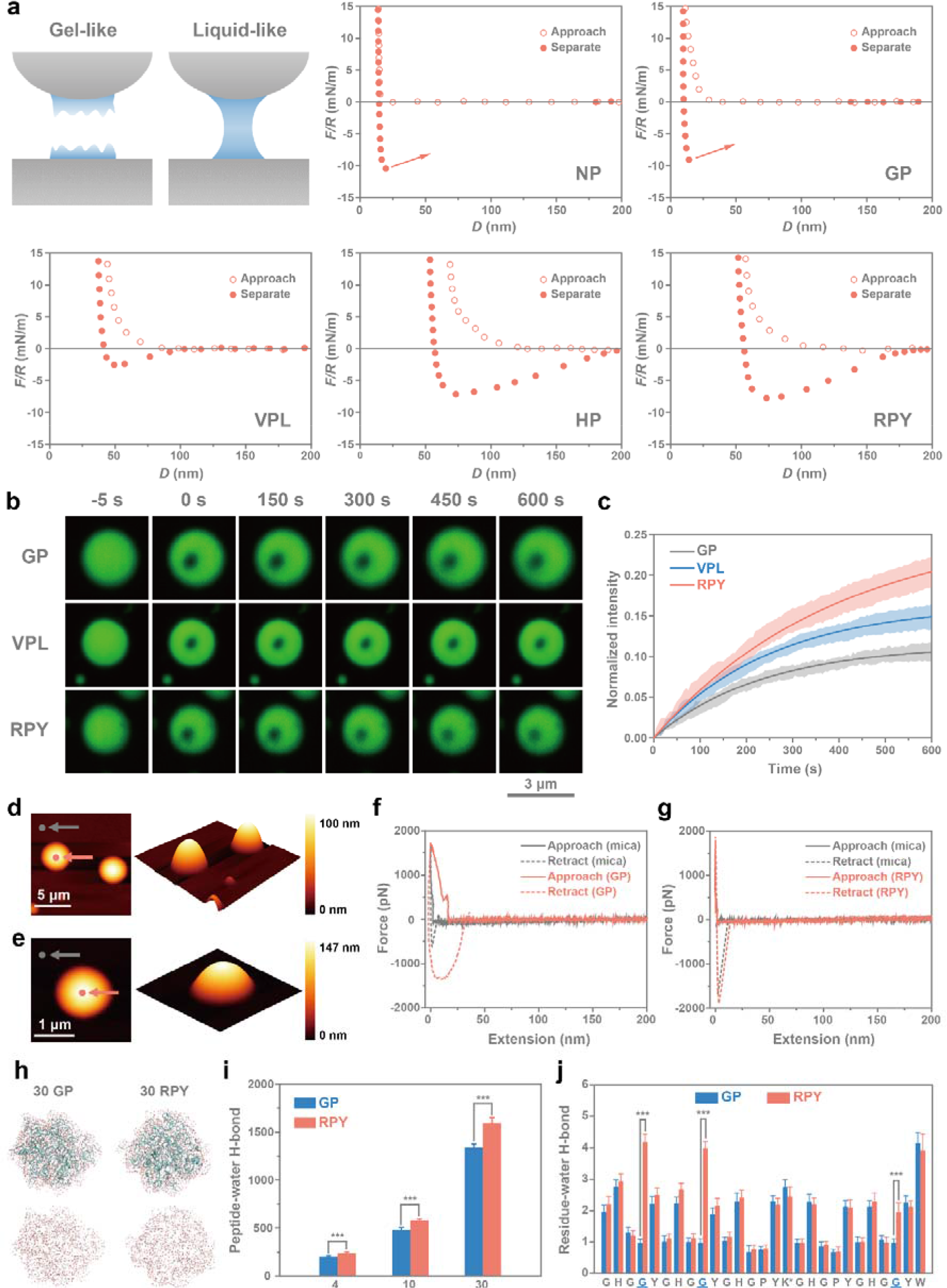
Characterization of CMs prepared from HB*pep*-SP variants. (a) Illustration of SFA measurements and force-distance (*F*-*D*) curves between cross-cylinders of CM variants, with *F* normalized by the radius of the cylindrically curved surfaces. **(b-c)** Fluorescence micrographs (b) and recovery curves (c) of CM variants. Data are presented as the mean_±_SD of *n* = 3 independent experiments, the errors are shadowed. **(d-g)** AFM nanoindentation measurements using a cantilever with stiffness 0.2 N/m. 2D and 3D images of GP (d) and RPY (e) CMs, with the location of force measurements on the CMs and the mica substrate indicated with red and grey arrows. Force traces for GP (f) and RPY (g) CMs compared to mica. (**h-j**) MD simulations of CM variants showing snapshots (last 300 ns of 1 μs simulations) of clusters formed by 30 GP and RPY peptides withholding water molecules (h); total number of hydrogen bonds between peptide and water for different cluster sizes (i), and average number of hydrogen bonds between individual residues and water molecules in each 30 peptides cluster (j). Data are presented as the mean_±_SD of *n* = 9000 independent frames; two-sided Student’s t-test, \**P*_<_0.05, \*\**P*_<_0.01, \*\*\**P*<0.001.

Fluorescence recovery after photobleaching (FRAP) was used to further explore the molecular mobility within single CMs,(*26, 27*) which was carried out by mixing fluorescently-labelled HB*pep*-SP variants with unlabeled peptides in a 0.5:99.5 ratio to minimize the influence of the fluorophore on the phase behavior. GP CMs exhibited the slowest recovery rate among the three tested CMs (Figs. 2b-c), indicating a reduced molecular mobility, which is consistent with SFA measurements that denoted gel-like properties. Conversely, RPY CMs displayed higher fluidity with the fastest recovery rate.

To substantiate the differences in viscoelastic properties, we carried out AFM nanoindentation measurements of CMs (Methods). For GP CMs, on the approach cycle, the force increased in the contact region with a slope that was significantly shallower compared with the force curve on the solid substrate (Figs. 2d and f). Furthermore, the retraction cycle showed hysteresis due to energy dissipation processes inside the CMs. Using the Sneddon model,(*28, 29*) the average elastic moduli of GP CMs was 2.4 ± 1.4 MPa for droplets deep enough to alleviate substrate effects (see Supplementary Note 1). On the contrary, for RPY CMs the force traces were essentially identical on the droplets and the solid substrate, indicating that the AFM tip penetrated through RPY CMs without resistance, until it reached the substrate whereupon a contact force was detected (Figs. 2e and g), an observation that corroborates the fluidic nature of RPY CMs.

To understand the differences in viscoelastic response at the molecular level, we conducted independent molecular dynamics (MD) simulations of 4, 10 and 30 chains of GP and RPY variants in aqueous conditions for 1000 ns (Methods). At equilibrium, both GP and RPY CMs contained large amounts of water molecules in their clusters, however, a significantly higher number of peptide-water hydrogen bonds were seen in RPY, mainly arising from R4, R9, and Y25, compared to G4, G9, and G25 in GP clusters (Figs. 2h-j). Consequently, we attribute the fluidic-like behavior of RPY CMs compared to GP CMs to their higher molecular hydration (see Supplementary Note 2).

Residue mutations also influenced the cargo recruitment efficiency within self-assembled CMs. Compared to VPL and RPY CMs, the recruitment efficiency of GP and NP decreased for both enhanced green fluorescent protein (EGFP) and R-phycoerythrin (R-PE) (Fig. S3a). Using fluorescence-activated cell sorting (FACS), we quantified the EGFP cargo content within individual CMs and found that VPL and RPY exhibited *ca.* 3- and 5- fold higher EGFP intensity than GP (Fig. S4), possibly due to the lower peptide/EGFP intermolecular interactions achievable by Gly residues.

### Cellular uptake and release kinetics of HB*pep*-SP CM variants determined by their material properties

Previous studies have revealed that cellular internalization of drug delivery carriers depends on their mechanical properties.(*30, 31*) Since we could regulate the material properties of the CMs, we postulated that such variations could be exploited to control interactions with the membrane, and in turn cellular uptake efficacy. Giant unilamellar vesicles (GUVs) prepared from 1-palmitoyl-2-oleoyl-sn-glycero-3-phosphocholine (POPC) were utilized as a surrogate model bilayer to examine mechanical interactions between lipid bilayer and CMs. Interactions between GUV and GP CMs led to bending of the GUV’s lipid bilayer (Fig. 3a), likely induced by the higher rigidity of GP CMs, which may lead to increased energy barrier during internalization and consequently decreased uptake rates(*30*). Conversely, the more fluidic RPY CMs were deformed upon interaction with GUVs and wetted the lipid bilayer.

**Fig. 3.**
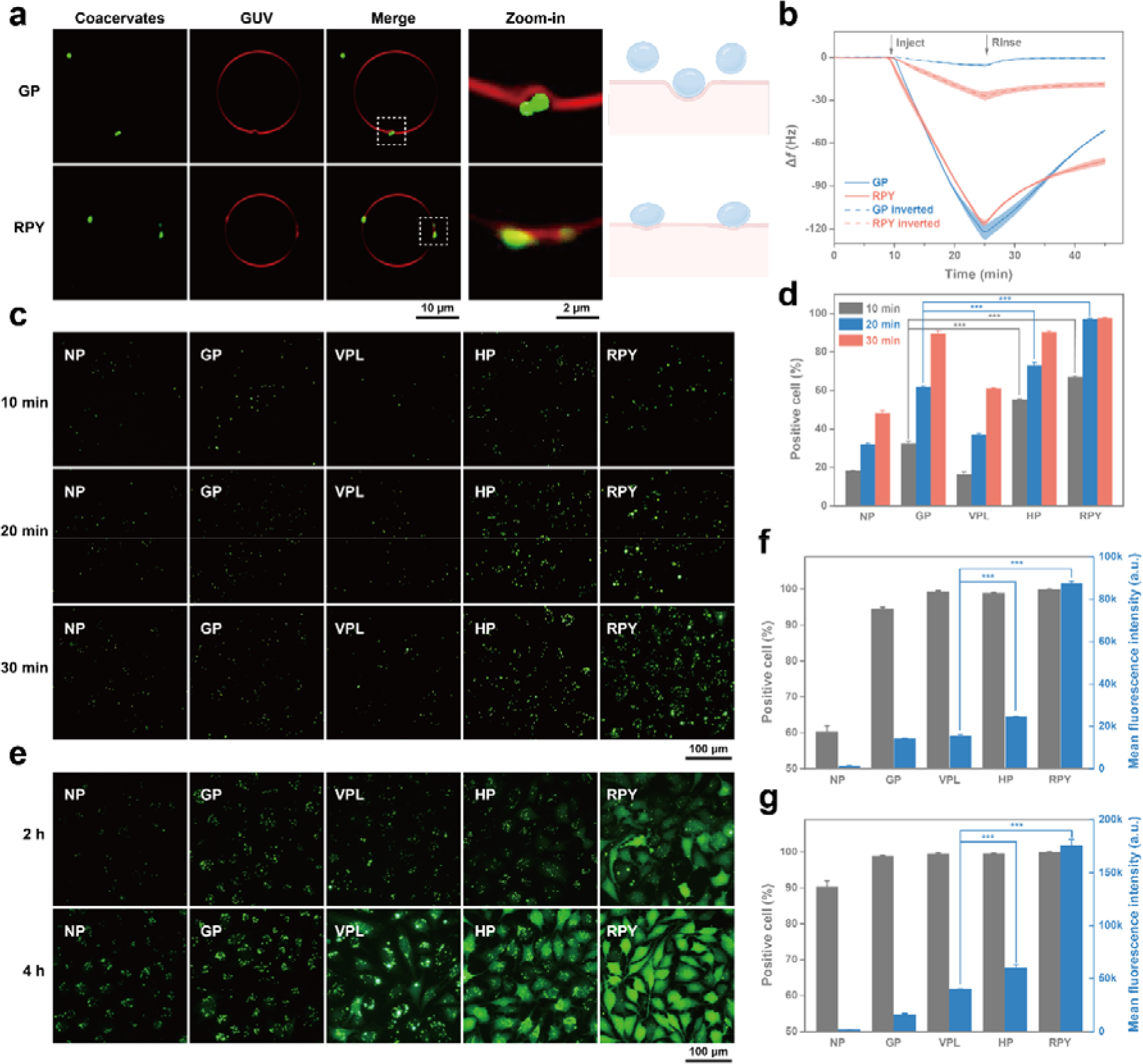
Membrane interaction, early internalization, and cargo release of CMs variants. (a) Fluorescence micrographs of GP and RPY CMs interacting with GUVs, illustrating GUVs’ bilayer bending by the stiffer gel-like GP CMs (top), and deformation and wetting of RPY CMs on GUVs’ bilayer (bottom). **(b)** QCM frequency shifts (Δ*f*) as a function of time for GP and RPY CMs adsorbed on supported lipid bilayers in the normal and inverted modes. Data are presented as the mean_±_SD of *n* = 3 independent experiments, the errors are shadowed. **(c-d)** Fluorescence micrographs (c) and FACS measurements (d) of HeLa cells treated with EGFP-loaded CMs formed by HB*pep*-SP variants for 10, 20 and 30 minutes. **(e)** Fluorescence micrographs and **(f-g)** FACS measurements of HeLa cells treated with EGFP-loaded CMs formed by HB*pep*-SP variants for 2 (f) and 4 hours (g). Data are presented as the mean_±_SD of *n*_ = 3 independent experiments; two-sided Student’s t-test, \**P* _< _0.05, \*\**P* _< _0.01, \*\*\**P* < 0.001.

Next, we assessed adhesive interactions between CMs and lipid bilayer membranes by carrying out quartz crystal microbalance (QCM) measurements of CMs on supported lipid bilayers (SLBs). In the normal mode, wherein the SLB is situated at the bottom of the flow cell, both GP and RPY CMs displayed high initial binding as evidenced by the decreased frequency (*f*), followed by rapid recovery upon rinsing. Compared to GP, RPY CMs were more resistant to wash-off during the rinsing step, suggesting stronger membrane binding (Fig. 3b). In order to alleviate the effect of gravitational sedimentation from the dense CMs, measurements were also carried out in the inverted mode,(*32*) whereby the flow cell was flipped upside down and SLBs oriented in the downward direction. RPY CMs showed significantly higher interaction with SLBs than GP CMs, persistently adhering to the bilayer even after 20 min of rinsing and causing a Δ*f* of – 21 Hz.

We then undertook cell uptake studies of CM variants in HeLa cells. HeLa cells treated with more viscous CMs (HP and RPY) loaded with EGFP (Figs. 3c-d) displayed significantly higher rates of EGFP-positive cells compared to gel-like GP and NP CMs after short incubation times (10 and 20 min), with RPY CMs exhibiting nearly 100% uptake after 20 min compared to 70%, 60% and 30% for HP, GP, and NP CMs, respectively. Given their similar size, this enhanced uptake of RPY CMs is attributed to their positive zeta potential arising from the Arg residue at the X1 and X2 positions (Fig. S5), facilitating adhesive interactions with the negatively charged cellular membrane. Despite VPL CMs displaying liquid-like properties, they exhibited lower uptake rates than GP CMs, which we attribute to incomplete phase separation caused by their more hydrophobic nature. Indeed, fewer CMs were formed at pH 6.5 as evidenced by half the count of CMs for VPL compared to GP and RPY in FACS measurements (Fig. S4b).

CMs disassembly (and concomitant release of cargos) is activated by GSH-induced disulfide bond reduction, which leads to cleavage of the sidechain grafted to the Lys residue of HB*peps*(*17*). Thus, the disassembly kinetics can be monitored by measuring the concentration decay of non-reduced HB*pep*-SP*s* upon incubation with GSH by high-performance liquid chromatography (HPLC). This decay could be adequately fitted by a first-order kinetic model and occurred faster in RPY CMs compared to GP CMs (Fig. S6), with a 1.8-fold higher rate constant. Since the grafted moiety is identical for both peptides, we attribute this disparity to the internal structure and material properties of the CMs. Owing to their higher molecular mobilities, we reason that the liquid-like RPY CMs enabled faster diffusion of GSH, thus accelerating the chemical reduction and initiating faster CMs disassembly. This was further substantiated by fluorescence microscopy and FACS measurements of HeLa cells treated with EGFP-loaded CMs. The faster release of EGFP from VPL, HP and RPY CMs after 2 and 4 h of incubation was evident from the homogenous fluorescence signal fulfilling the entire cells (particularly striking for HP and RPY CMs) as opposed to discrete puncta for cells treated with NP and GP CMs, which is indicative of non-disassembled CMs (Figs. 3e-g). While most cells internalized CMs after 2 h regardless of the peptide variant, there was a clear correlation between peptide variant and release kinetics as assessed by the mean fluorescence intensities (MFI) that directly reflects EGFP release in the cytosol. In the more liquid-like CMs, release kinetics increased in the order RPY > HP > VPL.

### Intracellular delivery of proteins and peptides mediated by CM variants

A key implication of the above results is that the intracellular release profile of macromolecules may in principle be finely regulated by simple amino acid adjustments. To verify this hypothesis, the 24 h delivery efficacy for all peptide variants was systematically assessed for a broad range of modalities. In line with short incubation times, RPY-mediated delivery of EGFP (Fig. 4a) yielded the highest MFI, indicative of the best release efficacy. While HP variant demonstrated superior EGFP release after 2 and 4 h, its recruitment ability was inferior to VPL; hence no significant differences were observed for these variants (Fig. 4b). We note that all three variants performed better than the commercial reagent PULSin. Although NP, SP, GP, and AP CMs also achieved over 95% cellular uptake (Figs. 4b and S7), their lower recruitment efficiency and slower release rate led to much weaker MFI. Variants incorporating charged residues displayed higher threshold concentrations to induce LLPS, leading to premature disassembly and cargo release during dilution in cell culture media (Fig. S7). The notable exception was RPY CMs, likely stabilized by cation-π interactions between positively charged and aromatic residues (*e.g*., R, Y and W)(*33*), as seen in the MD simulations (see Supplementary Note 2).

**Fig. 4.**
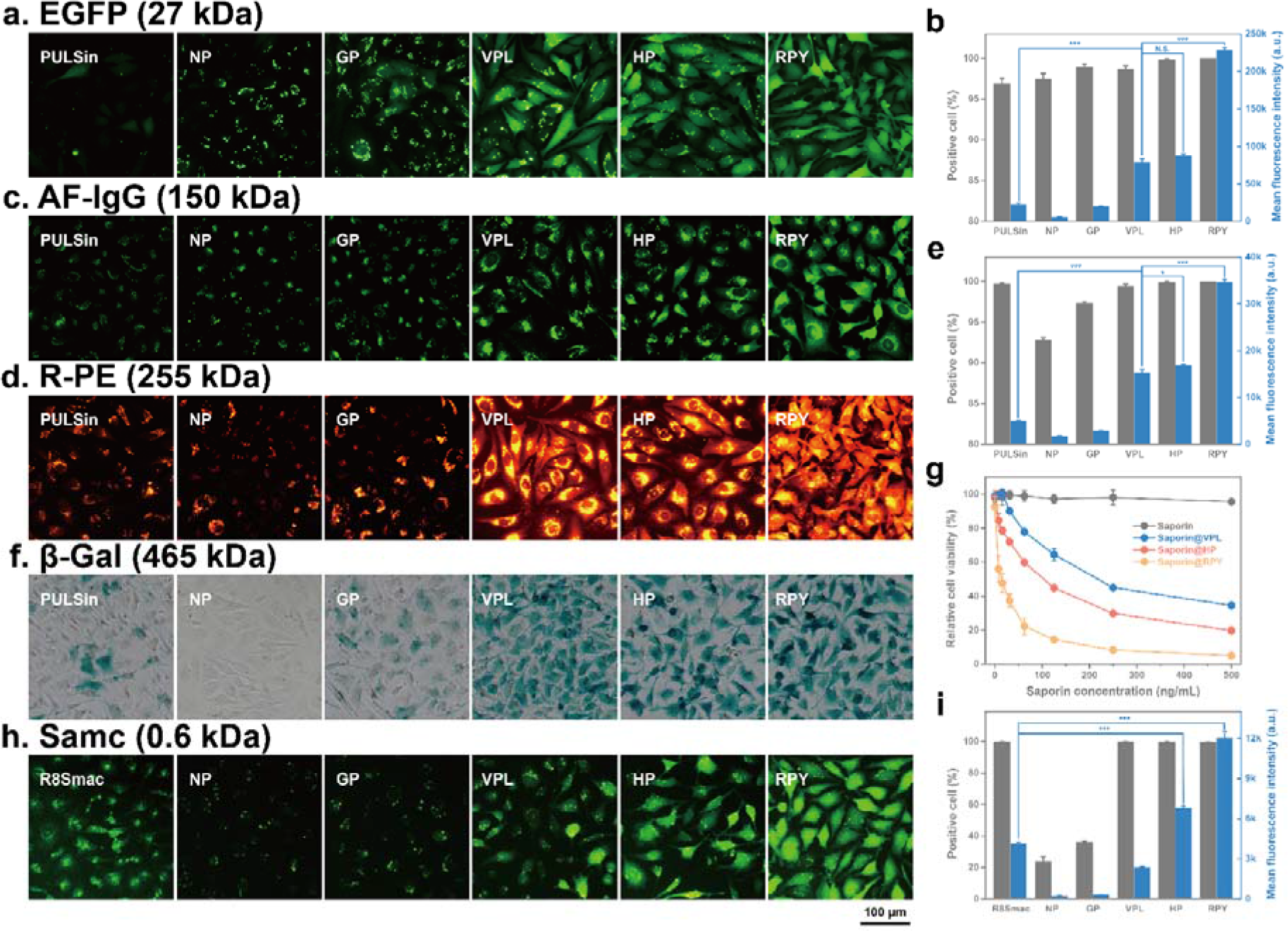
Intracellular delivery of proteins and peptides mediated by HB*pep*-SP CM variants. (a-b) Fluorescence micrographs (a) and FACS measurements (b) of HeLa cells treated with EGFP-loaded CMs variants for 24 hours compared to the commercial reagent PULSin. **(c)** Fluorescence micrographs of HeLa cells treated with AF-IgG-loaded CMs variants for 24 hours compared to PULSin. **(d-e)** Fluorescence micrographs (d) and FACS measurements (e) of HeLa cells treated with R-PE-loaded CMs variants for 24 hours compared to PULSin. **(f)** X-Gal staining of cells treated with β-Gal-loaded CMs variants for 24_hours compared to PULSin. **(g)** Concentration-dependent cytotoxicity of free saporin and saporin-loaded CMs variants. **(h-i)** Fluorescence micrographs (h) and FACS measurements (i) of HeLa cells treated with FITC-Smac-loaded CMs variants for 24 hours compared to Smac fused to the cell-penetrating peptide R8 (FITC-R8Smac) at the same concentration of 50 μM. Data are presented as the mean_±_SD of *n* = 3 independent experiments; two-sided Student’s t-test, \**P*_ < 0.05, \*\**P*_ < 0.01, \*\*\**P* < 0.001.

We then explored the transfection efficacy of proteins with a large spectrum of molecular weights (MWs), including the cytotoxic protein saporin (28.6 kDa), the Alexa Fluor 488-labeled immunoglobulin G antibody (AF-IgG, 150 kDa), R-PE (255 kDa), and β-galactosidase (β-Gal, 465 kDa) (Figs. 4c-g). In all cases, the transfection efficiency as quantified by the MFI (AF-IgG and RP), cell death (saporin), or enzymatic activity (β-Gal) followed the same performance trend of RPY > HP > VPL > GP > NP. For example, free saporin failed to enter HeLa cells and remained harmless (Fig. 4g), whereas VPL CMs-mediated delivery of saporin induced 65% cell death, which increased to 80% and 95% for HP and RPY CMs, respectively.

A similar trend was observed when delivering the second mitochondria-derived activator (Smac) short peptide (sequence: AVPIAQK), a potent apoptosis initiator.(*34*) Here, the transfection efficiency of the best-performing CMs (RPY) showed 3-fold higher MFI compared to Smac fused to the cell-penetrating peptide (CPP) octa-arginine (R8Smac), considered to be a highly efficient *in vitro* peptide delivery vector (Figs. 4h-i).(*35*)

### Gene transfection mediated by HB*pep*-SP CM variants

Gene therapy has been extensively investigated to treat various diseases,(*36, 37*) making significant contributions to the development of therapeutics, including clustered regularly interspaced short palindromic repeats (CRISPR) gene editing tools,(*38, 39*) COVID-19 vaccines,(*40, 41*) and chimeric antigen receptor (CAR) T-cell therapies.(*42, 43*) We have previously demonstrated that VPL CMs could deliver mRNA and all CRISPR/Cas9 modalities with good efficiency.(*12, 17*) We decided to expand the selection of nucleic acid therapeutics and explored the transfection efficacies of CM variants. In contrast to its low efficiency for protein and peptide delivery, GP CMs exhibited robust EGFP-encoding pDNA transfection in HeLa cells, significantly surpassing VPL CMs and Lipofectamine 2000 (Figs. 5a-b). This trend was further confirmed in the delivery of EGFP-encoding mRNA, where GP CMs transfected over 95% of cells with a 1.4-fold higher MFI compared to VPL (Figs. 5c-d). Similarly, HP and RPY variants demonstrated excellent efficiencies, transfecting 91.3% and 94.3% of cells for pDNA, and 99.5% and 98.7% for mRNA, respectively (Figs. 5a-d). The increased efficiency for GP compared to VPL in delivering nucleic acids can be attributed to the higher peptide concentration required to achieve full cargo recruitment for GP (Fig. S3b), which indicates weaker peptide-nucleic acid interactions, in turn leading to facilitated release.

**Fig. 5.**
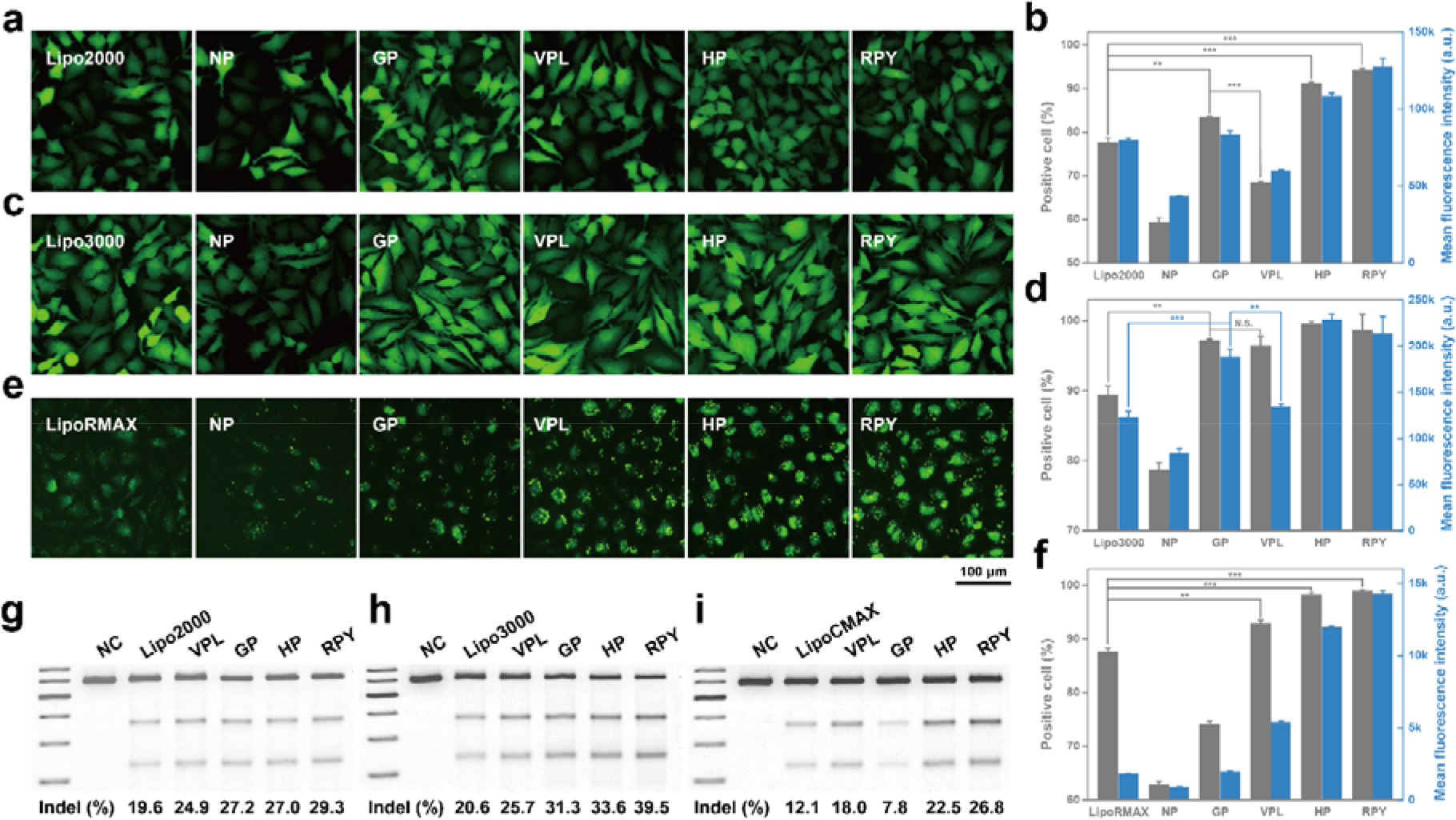
Gene transfection mediated by the HB*pep*-SP CM variant. (a-b) Fluorescence micrographs (a) and FACS measurements (b) of HeLa cells treated with EGFP pDNA-loaded CMs variants for 24 hours compared to Lipofectamine 2000 (Lipo2000). **(c-d)** Fluorescence micrographs (c) and FACS measurements (d) of HeLa cells treated with EGFP mRNA-loaded CMs variants for 24 hours compared to Lipofectamine 3000 (Lipo3000). **(e-f)** Fluorescence micrographs (e) and FACS measurements (f) of HeLa cells treated with FITC-siRNA-loaded CMs variants for 24 hours compared to Lipofectamine RNAiMAX (LipoRMAX). **(g-i)** Analysis of indel frequency at the HBB locus in HeLa cells treated with HBB-targeted CRISPR modalities-loaded CMs variants, including all-in-one pDNA compared to Lipo2000 (g), mRNA/sgRNA compared to Lipo3000 (h), and the RNP complex compared to Lipofectamine CRISPRMAX (LipoCMAX) (i). Data are presented as the mean_±_SD of *n* = 3 independent experiments; two-sided Student’s t-test, \**P* < 0.05, \*\**P* < 0.01, \*\*\**P*<0.001.

siRNA is another class of promising therapeutics that target and degrade mRNAs to prevent the expression of harmful proteins.(*44*). Given that the MW and working concentrations of siRNA were comparable to proteins, the transfection efficiency of the CMs variants exhibited a similar trend to protein delivery. Cells treated with CMs displayed increasing rates of positive cells and MFI in the order RPY > HP >VPL > GP >NP (Figs. 5e-f), with VPL, HP, and RPY CMs outperforming the highly specialized lipofectamine RNAiMAX (LipoRMAX). Using quantitative reverse transcription polymerase chain reaction (RT-qPCR) to quantify mRNA degradation induced by anti-PCSK9 siRNA, the siRNA-loaded HP and RPY CMs achieved mRNA knockdown rates of 74.2% and 86.5%, surpassing 50.3% mediated by LipoRMAX (Fig. S8). Furthermore, delivering anti-EGFP siRNA into HeLa cells expressing EGFP provided insights into the knockdown efficiency at the protein level. siRNA transfection mediated by HP and RPY CMs decreased the EGFP signal by 73.2% and 79.6%, respectively, significantly higher than LipoRMAX-mediated transfection (Fig. S9). For the VPL CM variant, despite exhibiting adequate cellular uptake (Fig. 5f), its knockdown efficiency at both the mRNA and protein levels weas inferior to LipoRMAX (Figs. S8-9), corroborating its slower release kinetics.

Finally, we attempted to deliver all three different types of CRISPR/Cas9 gene editing tools. Using the T7 Endonuclease I (T7EI) assay to assess the editing efficiencies on the HBB locus, GP, HP, and RPY CMs exhibited higher insertion-deletion (indel) frequencies compared to VPL and lipofectamines in delivering the all-in-one pDNA and mRNA/sgRNA gene editing machineries (Figs. 4g-i). While GP displayed lower efficiency when delivering CRISPR/Cas9 using the ribonucleoprotein (RNP) complex, HP and RPY maintained their high efficacy, surpassing the highly specialized lipofectamine CRISPRMAX. The excellent knockout efficacy of the CMs delivery system was further illustrated by transfecting mRNA and EGFP-targeted sgRNA into HeLa-EGFP cells, with all three CMs reducing the EGFP signal more efficiently than Lipo3000 (Fig. S10).

### Intracellular delivery into hard-to-transfect cells mediated by HB*pep*-SP CM variants

Expanding beyond HeLa cells, we finally investigated the ability of fluidic-like CMs to deliver therapeutics into hard-to-transfect cells such as primary and immune cells, starting with primary human foreskin fibroblasts (HFF). All three CMs demonstrated increased cellular uptake of EGFP and significantly higher MFI compared to PULSin (Fig. 6a). Owing to their enhanced internalization and cargo release kinetics, HP and RPY again outperformed VPL CMs. This trend was similarly observed in mRNA delivery, where HP and RPY CMs transfected 44.9% and 67.1% of HFF cells, respectively, compared to 37.2% of cells for VPL and 32.7% for Lipo3000 (Fig. 6b). In Jurkat T-cells, HP and RPY CMs successfully delivered EGFP into up to 88.6% of treated cells, resulting in 2.8- and 4.3-fold higher MFI compared to PULSin (Fig. 6c). For mRNA, 47.7% and 62.9% of cells transfected by HP and RPY CMs showed EGFP positivity, surpassing the 19.1% and 30.4% achieved by Lipo3000 and VPL CMs (Fig. 6d). As a final assessment, we aimed to transfect the RAW264.7 macrophage cell line, known for its innate resistance to foreign material transfection.(*45*) Remarkably, nearly all cells treated with our EGFP-loaded CMs exhibited positive signals (Fig. 6e), while only half of the cells were positive in PULSin-mediated delivery. For mRNA transfection, HP and RPY also demonstrated excellent efficacy of above 90% for macrophages (Fig. 6f). Although VPL exhibited lower efficiency (67.0%), it still significantly outperformed Lipo3000 (8.0%).

**Fig. 6.**
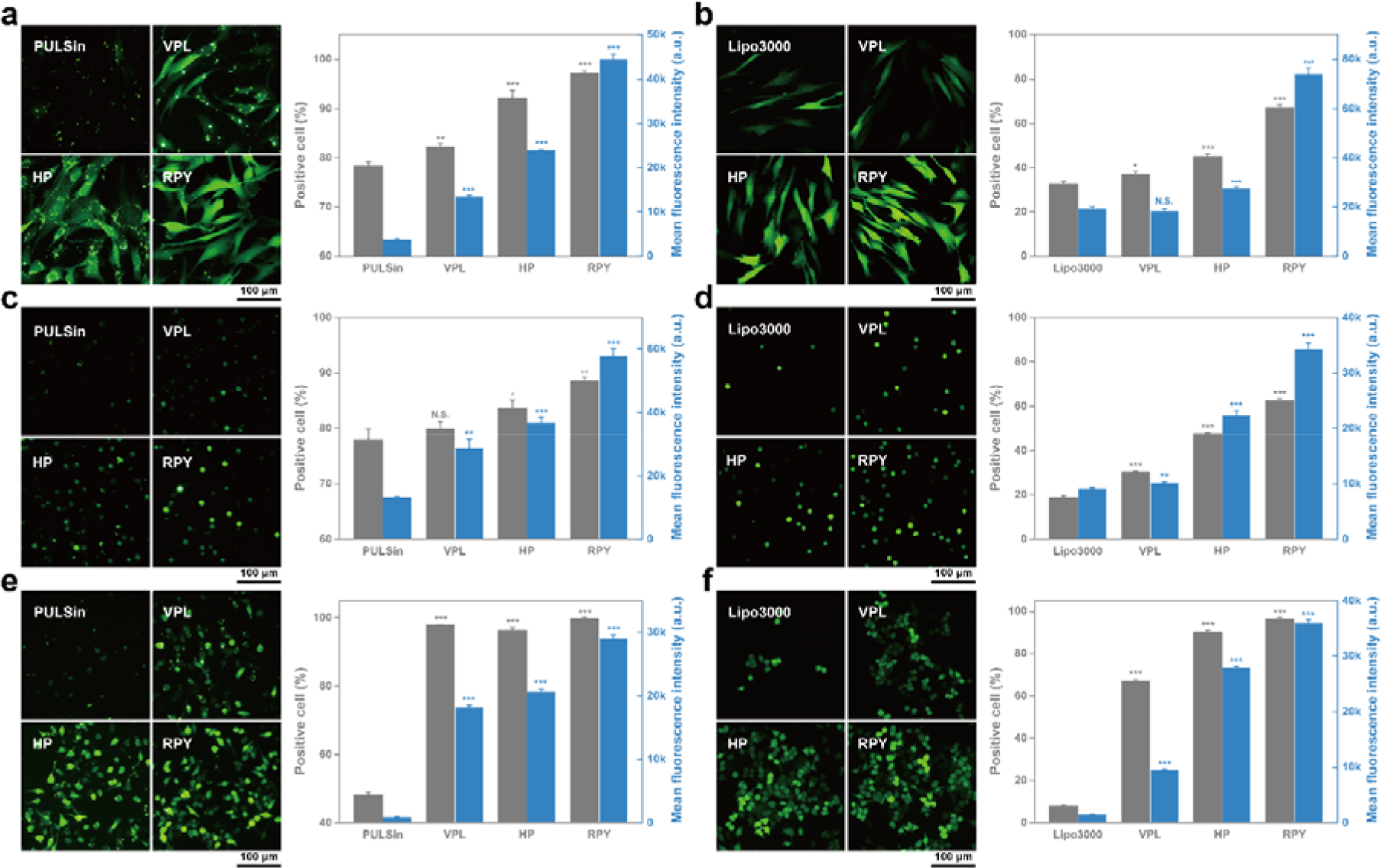
Intracellular delivery into hard-to transfect cells mediated by the HB*pep*-SP CM variants. (a,b) Fluorescence micrographs and FACS measurements of Human Foreskin Fibroblast (HFF) cells treated with EGFP-loaded CMs variants for 4 hours compared to PULSin (a), and treated with mRNA-loaded CMs variants for 24 hours compared to Lipo3000 (b). **(c,d)** Fluorescence micrographs and FACS measurements of Jurkat T-cells treated with EGFP-loaded CMs variants for 4 hours compared to PULSin (c), and treated with mRNA-loaded CMs variants for 24 hours compared to Lipo3000 (d). **(e,f)** Fluorescence micrographs and FACS measurements of macrophage RAW264.7 cells treated with EGFP-loaded CMs variants for 4 hours compared to PULSin (e), and treated with mRNA-loaded CMs variants for 24 hours compared to Lipo3000 (f). Data are presented as the mean_±_SD of *n* = 3 independent experiments; two-sided Student’s t-test, \**P* < 0.05, \*\**P* < 0.01, \*\*\**P* < 0.001 compared to commercial reagents.

## Discussion

Through systematic amino acid mutations, we have established molecular guidelines to design and optimize PSP CM-based intracellular delivery systems. Of particular significance is the inherent simplicity of the HB*pep*-SP sequence design comprising of tandem GHGXY repeats, which we have leveraged to develop a PSP toolbox comprising various HB*pep*-SP variants by systematically introducing all types of amino acid residues in the X positions. This versatile platform enables the delivery of an unprecedented broad spectrum of biomacromolecular therapeutics from a single delivery vehicle, including proteins, peptides, and nucleic acids across various cell types. Notably, PSP CMs exhibit superior efficiency compared to highly specialized delivery vehicles such as PULSin, lipofectamines, and CPPs, even demonstrating efficacy in notoriously hard-to-transfect cells such as primary and immune cells.

Beyond offering a robust intracellular delivery toolbox, our results establish the key material and biophysical properties of CMs that enable the control of uptake rates and intracellular release kinetics of cargos. Thus, fluidic-like CMs with enhanced adhesion and wetting to the cell membrane exhibit faster uptake and release kinetics of cargos compared to gel-like CMs. A key implication is that the release kinetics can in principle be modulated, from fast to slow release to meet a given therapeutic need. MD simulations suggest that peptide hydration plays an important role in differentiating between the fluidic versus gel-like properties of CMs, which can be modulated by the combination of specific amino acid residues at the X positions of the GHGXY repeats.

We believe that our findings may have broad practical implications, opening new avenues for the development of diverse therapeutic treatments encompassing areas such as vaccines, tissue engineering, and immunotherapies, while also providing a versatile research toolbox for the life sciences.

## Methods

### Materials

Resins and fluorenylmethoxycarbonyl (Fmoc)-protected amino acids used in solid-phase peptide synthesis were purchased from GL Biochem. N-hydroxysuccinimide, tetrahydrofuran, N,N′-diisopropylcarbodiimide, triphosgene and benzoic acid were purchased from Tokyo Chemical Industry (TCI). N,N-diisopropylethylamine, 2-hydroxyethyl disulfide, piperidine, trifluoroacetic acid, triisopropylsilane, saporin, β-galactosidase, R-phycoerythrin, Hoechst 33342, Oxyma, and reduced L-glutathione (GSH) were obtained from Sigma-Aldrich. Dichloromethane, N,N-dimethylformamide (DMF), Alexa Fluor 488 NHS ester, SYBR Safe DNA gel stain, Opti-MEM, 5-bromo-4-chloro-3-indolyl β-D-galactopyranoside (X-Gal), and Lipofectamine 2000, 3000, RNAiMAX and CRISPRMAX were purchased from Thermo Fisher Scientific. Cell counting kit-8 (CCK-8) was purchased from Abcam. PULSin protein transfection reagent was purchased from Polyplus. T7 Endonuclease I, Q5 Hot Start high-fidelity 2X master mix, and 100 bp DNA Ladder were purchased from New England Biolabs. Organic solvents, including ethyl acetate, hexane and diethyl ether were purchased from Aik Moh Paints & Chemicals Pte Ltd. Dulbecco’s modified Eagle medium (DMEM), RPMI-1640 medium, fetal bovine serum (FBS), phosphate-buffered saline (PBS) and antibiotic-antimycotic (100X) liquid were purchased from Gibco. EGFP-encoding pDNA, EGFP-encoding mRNA, Cas9 all-in-one pDNA, Cas9-encoding mRNA, HBB targeting sgRNA and EGFP targeting sgRNA were obtained from GenScript. FAM-siRNA (anti-PCSK9) was purchased from NAS Bioscience, and anti-EGFP siRNA was purchased from Integrated DNA Technologies. HeLa, Jurkat and RAW264.7 cell lines were obtained from ATCC. The stable EGFP expression HeLa cell line (HeLa-EGFP) was purchased from Cell Biolabs Inc. The primary human foreskin fibroblast (HFF) cells were a generous gift from Prof. Peter Dröge’s lab in the School of Biological Sciences, Nanyang Technological University, Singapore.

### Peptide synthesis and modification

The HB*pep*-SP variants were prepared by modifying the Lys side chain of the N-terminus protected Fmoc-HB*pep*-K peptide, followed by the Fmoc deprotection. First, the synthesis of Fmoc-HB*pep*-K backbone was conducted on a microwave-assisted solid phase peptide synthesizer (Liberty Blue) using N,N′-diisopropylcarbodiimide (DIC)/Oxyma as coupling reagents and 20% piperidine in DMF as deprotection reagents. After the synthesis, peptides were cleaved from the resins using a cocktail containing 95% of trifluoroacetic acid (TFA), 2.5% of H_2_O and 2.5% of triisopropylsilane (TIPS) for 2 h. Then the supernatants were collected by filtration and concentrated using a nitrogen flow, followed by precipitating into 50 mL of cold diethyl ether. The pellets from centrifugation were dried under vacuum and re-dissolved using 5% acetic acid aqueous solution for purification by HPLC (1260 Infinity, Agilent Technologies) equipped with a C8 column (Zorbax 300SB-C8, Agilent Technologies). The purified Fmoc-HB*pep*-K variants were isolated by lyophilization from HPLC elutes.

The Lys side chain amine of Fmoc-HB*pep*-K could react with an amine-reactive small molecule NHS-SS-Ph (Fig. S1), followed by deprotection to produce HB*pep*-SP as described in our previous works.(*12, 17*) In detail, 15 μmol of peptides was dissolved in 3 mL of dimethylformamide (DMF). Then, 450 μmol of N,N-diisopropylethylamine (DIEA) was added into the peptide solution followed by 100 μL of DMF containing 20 μmol of NHS-SS-Ph. After overnight reaction at room temperature, 1 mL of piperidine was added into the mixture for another 1 h of Fmoc deprotection. The raw products were precipitated out by adding 30 mL of cold diethyl ether and collected by centrifugation. The pellets were dried under vacuum and re-dissolved using 5% acetic acid aqueous solution for purification by HPLC. The purified HB*pep*-SP variants were isolated by lyophilization from HPLC elutes, which was confirmed by MALDI-ToF spectroscopy (AXIMA Performance spectrometer, Shimadzu), shown in Fig. S18.

### LLPS of HB*pep*-SP variants

The phase diagram of variants was determined by observing the phase separation of variants at different pHs and concentrations via an inverted microscope (AxioObserver.Z1, Zeiss). HB*pep*-SP variants were dissolved in 10 mM acetic acid aqueous solution with various concentrations as stock solutions. The LLPS was induced by mixing the stock solutions with buffers at a volume ratio of 1:9. The buffer details are described in Table S1.

### Surface force apparatus (SFA) measurements

The viscoelastic property of CM variants was measured using an SFA 2000 (SurForce LLC, Santa Barbara).(*23*) As described in previous studies,(*24, 46*) freshly cleaved mica with a 55 nm silver layer deposited on its back was glued on the glass disks. Then 20 μL of freshly prepared CMs (0.3 mM in PBS) was injected in the gap between two mica surfaces. The distance *D* between two surfaces was measured and calculated based on the fringes of equal chromatic order (FECO) technique. After the sample was injected into the gap, the system was equilibrated for 30 min by keeping two surfaces in contact with a bridging coacervate film. Then, the two surfaces started to approach followed by separation. The measured force *F* was normalized by the effective radius of the surface *R*.

### Fluorescence recovery after photobleaching (FRAP) experiments

To perform FRAP measurements, HB*pep*-SP peptide variants were labeled with Alexa Fluor 488 NHS ester on the N-terminus. The labeled peptides were mixed with pristine ones at a ratio of 0.5:99.5 to prepare stock solutions with a final concentration of 3 mM. The CMs were prepared by mixing the stock solution with the pH 6.5 buffer (VPL) or pH 7.0 buffer (GP and RPY) at a ratio of 1:9. The experiment started by applying a 488 nm laser pulse at the power of 50% on the chosen area within a single CM from the confocal microscope (Eclipse Ti2, Nikon) to bleach its fluorescence. The confocal microscope then took images of the sample every 5 s. The fluorescence intensity was quantified by using ImageJ software and normalized by the intensity before the photobleaching.

### Atomic force microscopy (AFM) measurements

The schematic of AFM measurements is shown in Fig. S19. After an initial immobilization of CMs on mica (Ted Pella, Inc.) for a duration of 5 min and removal of unbound droplets via buffer (PBS) exchange, an AFM tip (SNL, Bruker) was brought in proximity to the CMs. Single CMs were localized by imaging in tapping mode (frequency about 30 kHz, amplitude setpoint about 90%). Generally, images of individual CMs were acquired at a scan rate of 0.2–0.4 Hz and a resolution of 64 points per line. After localization, “Point & Shoot” function (NanoScope, Bruker) was used to place the AFM tip on the apex of selected CMs before applying indentation using ramp mode. As a control, points on solid surface near selected CMs were also indented. The CMs (and the control point on solid) were indented up to a threshold force value of about 2 nN at a preset frequency of 0.5 Hz. Indented CMs were imaged again in tapping mode to detect signs of degradation, drift, or displacement due to tip lateral forces. The height of individual CMs before and after indentation were calculated by placing a trace line over the particles on images previously flattened using Gwyddion 2.47.(*47*) For elasticity evaluation, we proceeded as follows: deflection versus piezo displacement was converted to force versus tip–surface separation (or indentation distance) using protocols written in Igor Pro (Wavemetrics).(*48*) Fit to the force versus indentation distance curve in the tip-CM contact region was used to calculate modulus based on Sneddon model (NanoScope, Bruker). Correction to the modulus was applied using bottom effect cone correction (BECC).(*29*) Ten indentations per CM were used for the evaluations. Average modulus of GP CMs was calculated from at triplicate independent preparations. Additionally, GP and RPY CMs were investigated using softer tips (MLCT-Bio, Bruker). Prior to the measurements, the AFM tip was cleaned using UV-ozone (Novascan). Dimension FastScan (Bruker) was used in all the measurements. The calibrated tip parameters included stiffness (about 0.2 N/m for SNL and 0.04 N/m for MLCT-Bio) using the thermal method, and optical lever sensitivity using cantilever–mica hard contact (set point equal to 0.4 V).(*48*)

### Cargo recruitment and redox-responsivity evaluations

The recruitment efficiency of CM variants was evaluated by measuring the cargo concentration in the supernatant after centrifugating the cargo-loaded CMs at 15,000 rpm for 5 min. For fluorescence protein cargos like EGFP and R-PE, the peptide stocks (3 mM in 10 mM acetic acid aqueous solution) were mixed with the buffers (pH 6.5 buffer for VPL and pH 7.0 buffer for other variants) containing 0.1 mg/mL proteins as a volume ratio of 1:9 to induce LLPS and recruitment. The mixtures were incubated at room temperature for 5 min and then centrifugated. The concentration of unrecruited cargos in the supernatant was determined by their fluorescence intensity measured by a plated reader (Infinite M200 Pro, Tecan) using 488 nm/519 nm (EGFP) and 560 nm/590 nm (R-PE) for the excitation/emission wavelengths. The EGFP-loaded CMs were also measured by FACS (LSR Fortessa X20, BD Biosciences) for the EGFP intensity within the CMs. For the pDNA cargo (10 μg/mL), agarose gel electrophoresis was used to evaluate the unrecruited cargo in the supernatant at various HB*pep*-SP variant concentrations. The gels were stained with SYBR Safe and imaged using a gel documentation system (iBright CL1500, Thermo Fisher).

The difference in the reduction rate of self-immolative side chain of different HB*pep*-SP variants was evaluated by measuring the concentration decrease of unreacted GP and RPY peptides in the presence of 1 mM GSH. The freshly prepared GP and RPY CMs (100 μL, 0.3 mM) were diluted in 900 μL of PBS containing 1.11 mM of GSH. The mixtures were incubated at 37[°C for different time periods before adding 50 μL of acetic acid to dissolve all the unreacted peptides, and their concentrations were measured by HPLC.

### Molecular dynamics (MD) simulations

Molecular dynamics (MD) simulations were carried out for two peptides GP and RPY to understand how intermolecular interactions modulate the properties of CMs. For each peptide, systems containing 4, 10, and 30 peptide molecules were simulated, respectively. Each system was subject to 3 replicates of simulations, with each replicate running for 1 μs, resulting in a total simulation time of 18 μs. In each simulation, the required number of peptide molecules was randomly placed in a cubic box and solvated with water molecules. To be consistent with in vitro experimental conditions, 0.16 M NaCl was added to each system. Each system was initially subject to 500 steps using steep descent energy minimization. Subsequently, a 100 ps of MD simulation in the NVT ensemble was carried out, followed by production simulations in the NPT ensemble that proceeded in three stages. The first stage involved a 300 ns of simulation at 300 K, during which the peptide was observed to aggregate and form irregular clusters. In the second stage, a 200 ns simulated annealing simulation was applied to accelerate equilibration. Finally, a 500 ns production run at 300 K was carried out. The details of each system are summarized in Table S2. The number of hydrogen bonds, proximal radial distribution functions (pRDF),(*49, 50*) solvent accessible surface area (SASA), and the number of π-π and cation-π interaction pairs were calculated using the combination of the last 300 ns of the three replicates.

In all simulations, the peptides were modelled using the AMBER14sb force field and water was described by the TIP3P model.(*51, 52*) Parameters of the unnatural amino acid KSP were obtained using the antechamber module of AMBER 20 package.(*53*) Lennard-Jones and short-range electrostatic interactions were computed using a cutoff of 0.9 nm, while long-range electrostatic interactions were calculated using PME.(*54*) All simulations were carried out in the NPT ensemble with temperature and pressure maintained at 300 K and 1 bar except for the simulated annealing simulations, which were conducted at a temperature of 400 K. All simulations were carried out using GROMACS 2021 patched with Plumed-2.9.(*55, 56*)

### Interactions between lipid bilayers and CMs

The giant unilamellar vesicles (GUVs) were prepared from 99.5% of 1-palmitoyl-2-oleoyl-sn-glycero-3-phosphocholine (POPC) and 0.5% of 1,2-dipalmitoyl-sn-glycero-3-phosphoethanolamine-N-(lissamine rhodamine B sulfonyl) (ammonium salt) (Rhod-PE) using the gel-assisted method described in previous studies.(*57*) To investigate the interactions between GUVs and CM variants, 100 μL of freshly prepared GP or RPY CMs (0.3 mM peptide, containing 0.5% Alexa Fluor 488 labeled peptide) were diluted in 900 μL of PBS containing GUVs. The mixture was visualized using a confocal microscope (Eclipse Ti2, Nikon).

The adhesion of CMs on the supported lipid bilayers (SLBs) was evaluated by quartz crystal microbalance (QCM). First, the SLBs were deposited on the silica sensor by solvent exchange method.(*58*) Then, PBS flowed into the QCM chamber at a flow rate of 0.05 mL/min until Δ*f* < 2 Hz in 10 min. After the system reached equilibrium, the mixture of 100 μL freshly prepared GP or RPY CMs (0.3 mM) and 900 μL of PBS flowed in at the flow rate of 0.05 mL/min for 15 min, followed by 20 min of rinse with PBS.

### Cell cultures

HeLa, primary human foreskin fibroblast (HFF), and RAW 264.7 cells were cultured in DMEM supplemented with 10% FBS, 100 U/mL penicillin, and 100 μg/mL streptomycin under typical conditions (37 °C and 5% CO2). Jurkat cells were cultured in RPMI-1640 Medium supplemented with 10% FBS, 100 U/mL penicillin, and 100 μg/mL streptomycin. HeLa-EGFP cells were cultured in DMEM supplemented with 10% FBS, 100 U/mL penicillin, 100 μg/mL streptomycin, and 10 μg/mL blasticidin.

For HeLa, HFF, and HeLa-EGFP cells, the subculture started by detaching the cells with trypsin treatment, followed by centrifugation (1000 rpm, 5 min) to collect the cells. Then the pellets were resuspended with fresh media for subculture or experiments. The Jurkat cells are suspension cells, the subculture was conducted by dilution of cell culture in fresh media to achieve the required cell density. For RAW 264.7 cells, the cells were detached from the culture flask using a cell scraper (Corning) and collected by centrifugation (1000 rpm, 5 min). Then the pellets were resuspended with fresh media for subculture or experiments.

### Protein and peptide delivery

To perform protein and peptide deliveries, cells were suspended in 1.5 mL of full media and transferred into 35 cm^2^ culture dishes. When the confluency reached ∼ 60%, the medium was replaced with 900 μL of Opti-MEM, then 100 μL of freshly prepared CMs (0.3 mM HB*pep*-SP variants, 0.1 mg/mL proteins or 50 μM peptide) were prepared by adding the HB*pep*-SP stocks into cargos containing buffer (pH 6.5 buffer for VPL and pH 7.0 buffer for other variants), and added into the Opti-MEM. The treated cells were imaged at different time points including 10 min, 20 min, 30 min, 2 h, and 4 h using fluorescence microscopy (AxioObserver.Z1, Zeiss), the cellular uptake and cargo release were also quantified by FACS (LSR Fortessa X20, BD Biosciences). After 4 h of incubation, the CMs-containing medium was removed, and the cells were washed with PBS twice before being cultured in 1.5 mL of fresh full medium. After another 20 h, cells were imaged under fluorescence microscopy and analysed by FACS (LSR Fortessa X20, BD Biosciences) to evaluate the 24 h release efficiencies of CM variants. The commercially available protein transfection reagent PULSin (Polyplus) was used as a comparison according to protocols from the manufacturers. The peptide delivery efficiencies of CM variants were compared with octa-arginine conjugated Smac (R8-Smac) at the same concentration of 50 μM.

### Gene transfection

To evaluate the gene transfection efficiency of CM variants, the pDNA and mRNA encoding EGFP reporter gene were used as cargos. Before transfection, cells were incubated in 35 cm^2^ dishes until the confluency reached ∼ 60%. The medium was then replaced with 900 μL of Opti-MEM, followed by the addition of 100 μL of freshly prepared pDNA- or mRNA-loaded CMs (0.3 mM HB*pep*-SP variants, 10 μg/mL pDNA or mRNA). After 4 [h of incubation, the medium was removed, and the cells were washed with PBS twice before adding 1.5 mL of the full medium. Transfection was then continued for another 20 h before imaging the cells under a fluorescence microscope and testing the transfection efficiency by FACS.

Two siRNAs including FAM labeled anti-PCSK9 and unlabeled anti-EGFP siRNA were used to evaluate the delivery efficiency of CMs. HeLa or HeLa-EGFP cells were cultured in 35 cm^2^ dishes with full medium before the transfection. Then, the medium was replaced with 900 μL of Opti-MEM and 100 μL of freshly prepared siRNA-loaded CMs (0.3 mM HB*pep*-SP variants, 200 nM siRNA). After 4 h of uptake, the cells treated with FAM-siRNA were imaged by fluorescence microscopy and the delivery efficiency was quantified by FACS. On the other hand, to measure the knock-down efficiency, after 4 h of uptake, the CMs containing medium was removed. The cells were washed with PBS twice and then cultured in 1.5 mL of full medium for another 20 h. The PCSK9 mRNA knock-down efficiency was measured by reverse transcription-quantitative polymerase chain reaction analysis (RT-qPCR) and normalized by comparing it to glyceraldehyde 3-phosphate dehydrogenase (GADPH) mRNA. Meantime, by delivering anti-EGFP siRNA into HeLa-EGFP cells, the protein level knock-down efficiency could be visualized by fluorescence microscopy and quantified by FACS.

To deliver all three types of CRISPR/Cas9 genome editing modalities, HeLa cells are cultured in 35 cm^2^ dishes until reaching 40% confluence. Then the medium was replaced with 900 μL of Opti-MEM and 100 μL of cargo-loaded CMs. The final cargo concentration is 2 μg/mL for the all-in-one pDNA, 2 and 1 μg/mL for the mRNA and sgRNA mixture, and 2 and 1 μg/mL for the Cas9 nuclease and sgRNA complex. After 4 h of uptake, the medium was discarded. The cells were washed with PBS twice and cultured in full media for another 44 h. The efficiency of 48 h of transfection can be evaluated by using the T7 Endonuclease 1 (T7EI) assay. First, the genomic DNA was extracted by using a DNeasy blood and tissue kit (QIAGEN). The target genomic locus was amplified by PCR using Q5 Hot Start high-fidelity 2X master mix (NEB) and primers listed in Table S3 and purified by PureLink PCR purification kit (Thermo Fisher Scientific). Then, 200 ng of PCR products were digested by T7EI and analyzed by 2% agarose gels before imaging with the gel documentation system. The gray level of digested bands and undigested bands was measured by ImageJ. The indel percentage could be calculated by the following formula:(*59, 60*)

[1−(1−fraction cleaved)^1/2^]×100

where fraction cleaved = the sum of each digested band intensity/(the sum of each digested band intensity + undigested band intensity).

The knock-out efficiency was also quantified by delivering Cas9 mRNA and EGFP-targeting siRNA into HeLa-EGFP cells following prior protocols. After 48 h of transfection, treated HeLa-EGFP cells were imaged under a fluorescence microscope, and their EGFP intensity decrease was measured by FACS.

### Cytotoxicity study

The cytotoxicity of the saporin-loaded or pristine CMs was evaluated using the Cell Counting Kit-8 (CCK-8). As described previously,(*61*) cells were cultured in 96-well plates with 100 μL of full media and incubated for 24 [h. The medium was then replaced with 100 μL of Opti-MEM containing saporin-loaded CMs (various concentrations of saporin, 0.3 mM HB*pep*-SP variants) or various concentrations of HB*pep*-SP variants. After 4 h of uptake, the medium was removed, and the cells were washed with PBS twice and cultured in 100 μL of fresh full medium. The cells were incubated for another 20 h before changing the medium to the full medium containing 10% CCK-8 solution. After 4 h of incubation, the cells were measured for absorbance at 460 nm using a microplate reader (Infinite M200 Pro, Tecan). The relative cell viability was calculated as:

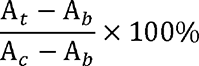

where *A*_t_, *A*_b_, and *A*_c_ represent the absorbance of tested cells, no cells, and untreated cells, respectively.

CM variants showed negligible cytotoxicity in all four cell lines including HeLa, HFF, Jurkat and RAW 264.7 (Fig. S20).

### Statistics and reproducibility

All experiments were repeated three times. The data are presented as mean[±[standard deviation (SD). Statistical significance (\**P* < 0.05, \*\**P* < 0.01, \*\*\**P* < 0.001) was evaluated using a two-sided Student’s t-test when two groups were compared. All microscopy experiments were repeated independently three times and the presented images are representative of the obtained data.

## Acknowledgements

This research was funded by the Singapore Ministry of Education (MOE) through an Academic Research Fund (AcRF) Tier 3 grant (grant no. MOE 2019-T3-1-012). We thank the A*Star for support and computing resources. J.Y. acknowledges support from the Singapore National Research Fellowship (NRF-NRFF11-2019-0004).

## Data availability

All data needed to evaluate the conclusions in the paper are present in the paper and/or the Supplementary Materials.

## Supplementary information

## Supplementary Note 1

### AFM measurements

Examples of AFM measurements with the GP CM are shown in Fig. 2f (cantilever stiffness 0.2 N/m) and in Fig. S11 (cantilever stiffness 0.04 N/m). With both cantilever stiffnesses, a viscoelastic response was detected, which was particularly significant in comparison with the force response on hard solid. Similar to GP CMs, RPY CMs were investigated using the two different AFM tips. Examples of the measurements with RPY CM are shown in Fig. 2g (cantilever stiffness 0.2 N/m) and in Fig. S12 (cantilever stiffness 0.04 N/m). With both cantilever stiffnesses, the force response was akin to the force response on hard solid which indicates that the tip penetrated RPY CMs with no resistance (*i.e.*, liquid-like behavior).

The distribution of elastic moduli of GP CM, varying from about tens of MPa to a few MPa, is shown in Fig. S13a. The reported data points present an average modulus from ten consecutive measurements of individual GP CMs. We did not observe a significant variation in the elastic moduli between the ten measurements. Correlation of the elastic moduli with the height profiles of individual GP CMs showed effects from the underlying substrate (despite BECC correction(*1*) or potentially due to a limited volume effect (Fig. S13b). Therefore, to report an effective elastic modulus, we averaged the elastic moduli of those GP CMs with a height above 70 nm. With this correction, the average elastic modulus of GP CM was 2.4 ± 1.4 MPa.

## Supplementary Note 2

### MD simulations

To characterize peptide hydration, we calculated the proximal radial distribution function (pRDF) for all clusters.(*2*) Notably, RPY displayed higher peaks in pRDF across all clusters (Fig. S14). Furthermore, RPY demonstrated a larger solvent-accessible surface area (SASA) compared to GP (Fig. S15). Both SASA and pRDF results suggest a greater degree of hydration for RPY relative to GP. The introduction of Arg enables RPY to engage in cation-π interactions with aromatic residues, which could play an important role in modulating the properties of the CM. As illustrated in Figs S16 and S17, RPY engages in many cation-π interactions arising from its R, Y and W residues, contributing to its adhesive properties.

---

**Fig. S1.**
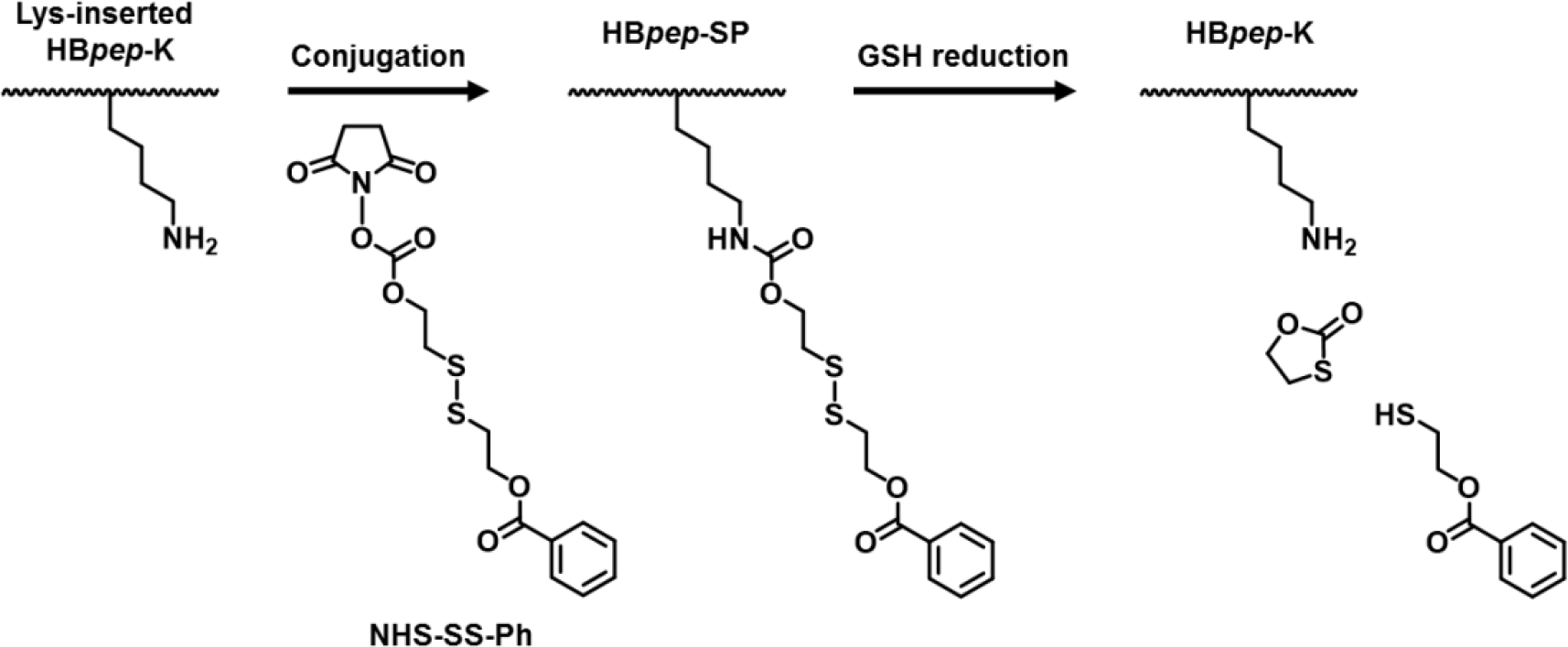
The conjugation of redox-responsive moiety on the Lys side chain of HB*pep*-K and its cleavage induced by GSH.

**Fig. S2.**
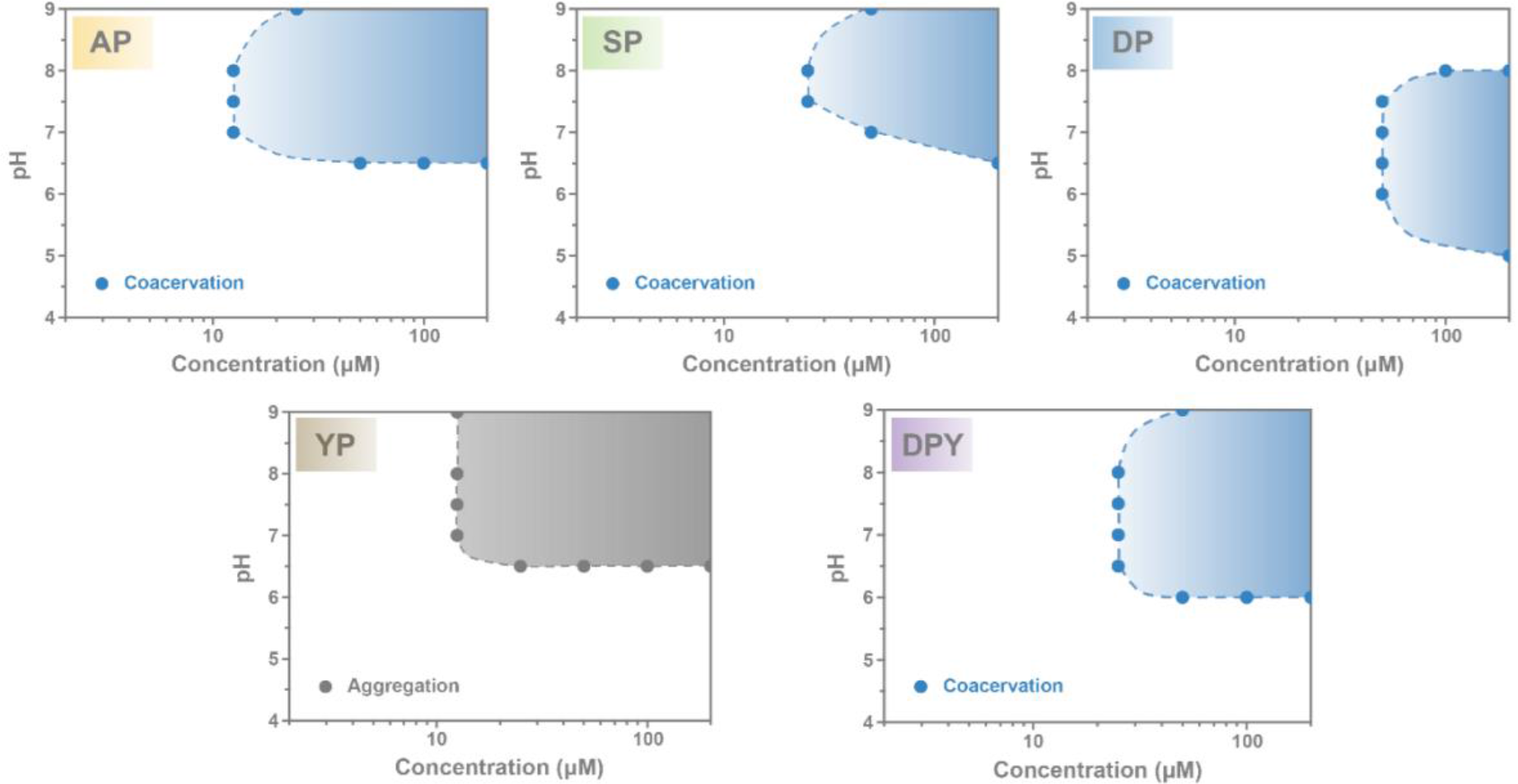
Phase diagram of representative HB*pep*-SP variants at the ionic strength of 100 mM, with the region of coacervation or aggregation shadowed in blue and grey, respectively.

**Fig. S3.**
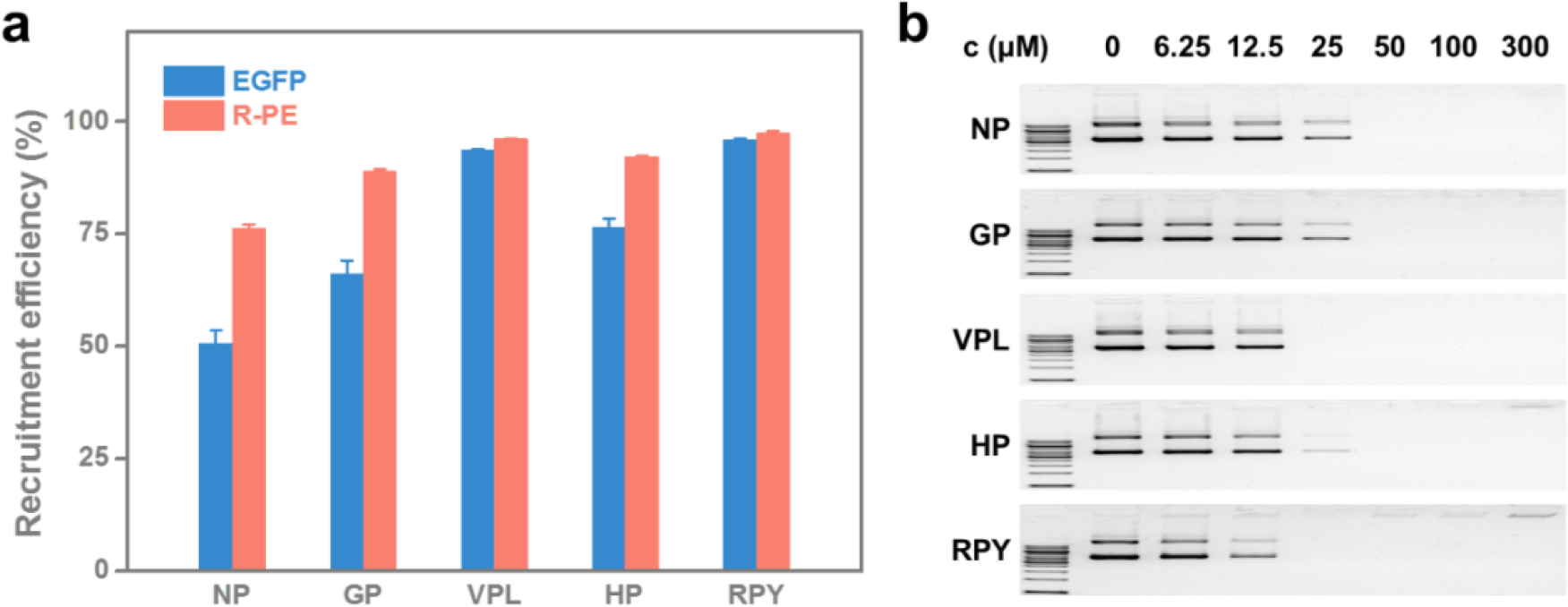
Recruitment efficiency of biomacromolecules by different CMs variants. **(a)** Recruitment efficiency of EGFP and R-PE (0.1 mg/mL) using variant CMs evaluated by fluorescence measurements on the supernatant after centrifugation. Data are presented as the mean ± SD of n = 3 independent experiments. **(b)** Recruitment efficiency of pDNA (10 μg/mL) using CMs variants at various concentrations evaluated by agarose gel electrophoresis on the supernatant after centrifugation. No band on the gel indicates full recruitment of cargos within the CMs.

**Fig. S4.**
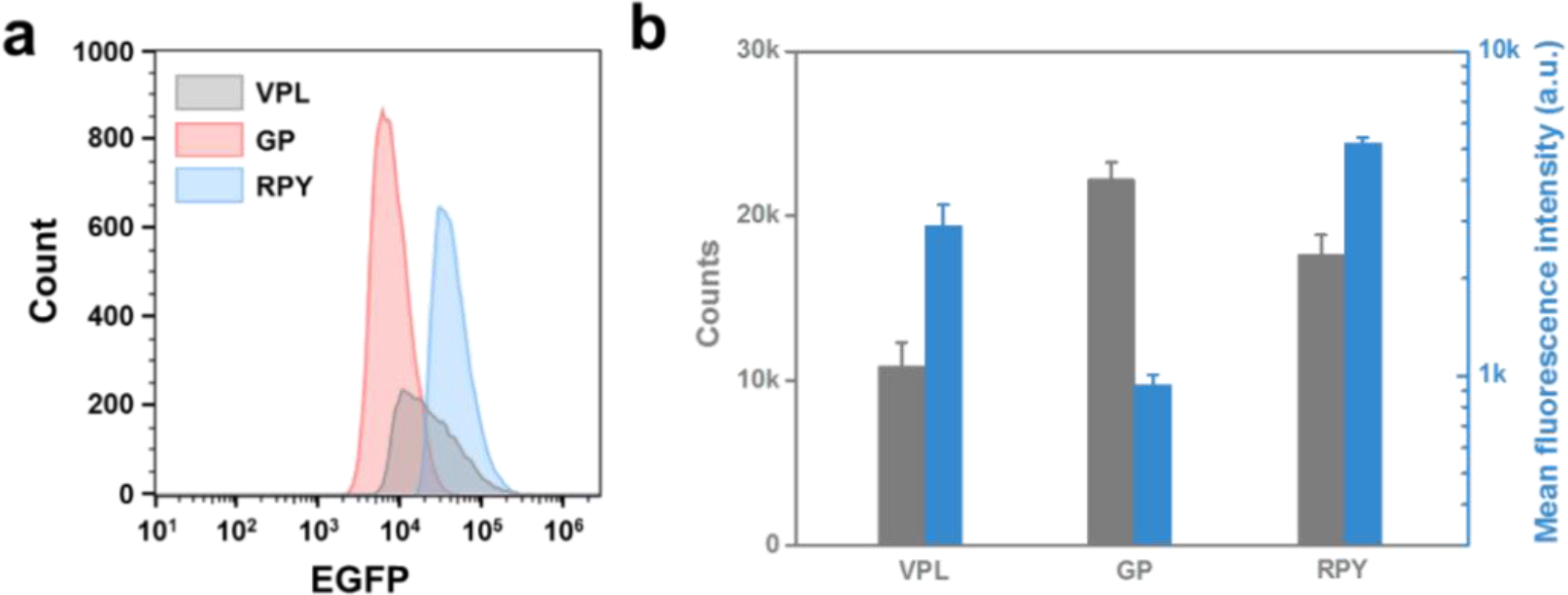
FACS measurements of EGFP-loaded CMs variants. **(a)** Representative histogram of the EGFP signals measured from EGFP-loaded VPL, GP and RPY CMs. **(b)** Average counts and mean fluorescence intensity of the EGFP-loaded CM variants. Data are presented as the mean ± SD of n = 3 independent experiments.

**Fig. S5.**
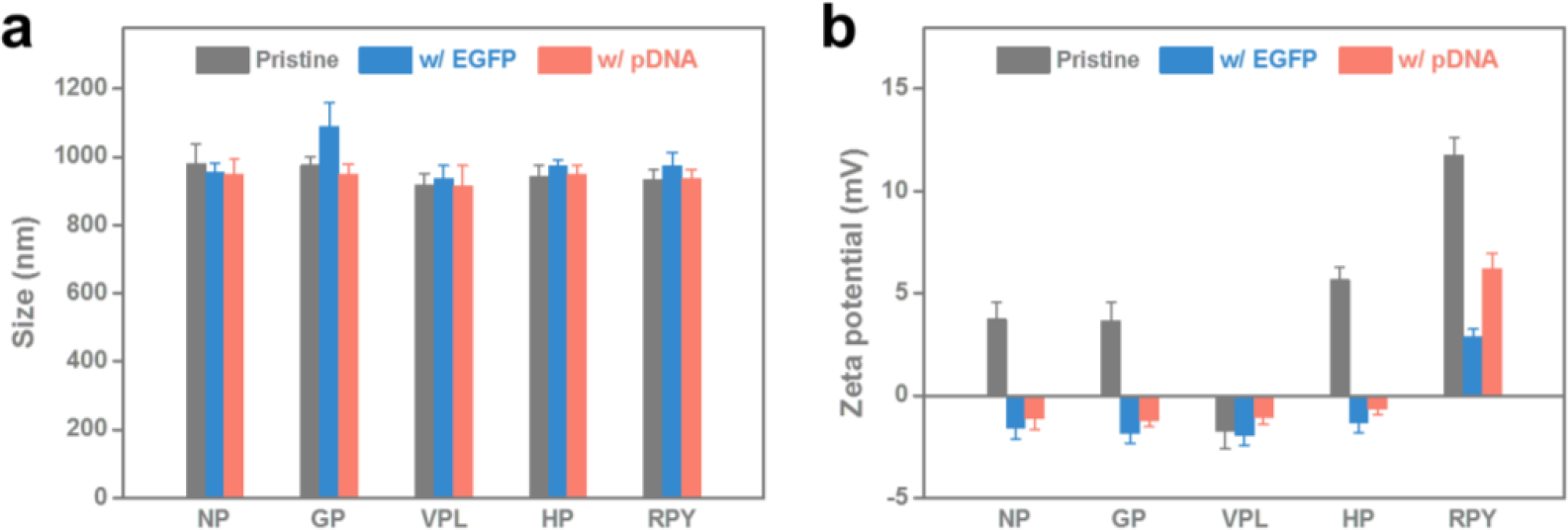
The Size **(a)** and zeta potential **(b)** of CM variants with and without cargos measured by DLS. Data are presented as the mean ± SD of *n* = 3 independent experiments.

**Fig. S6.**
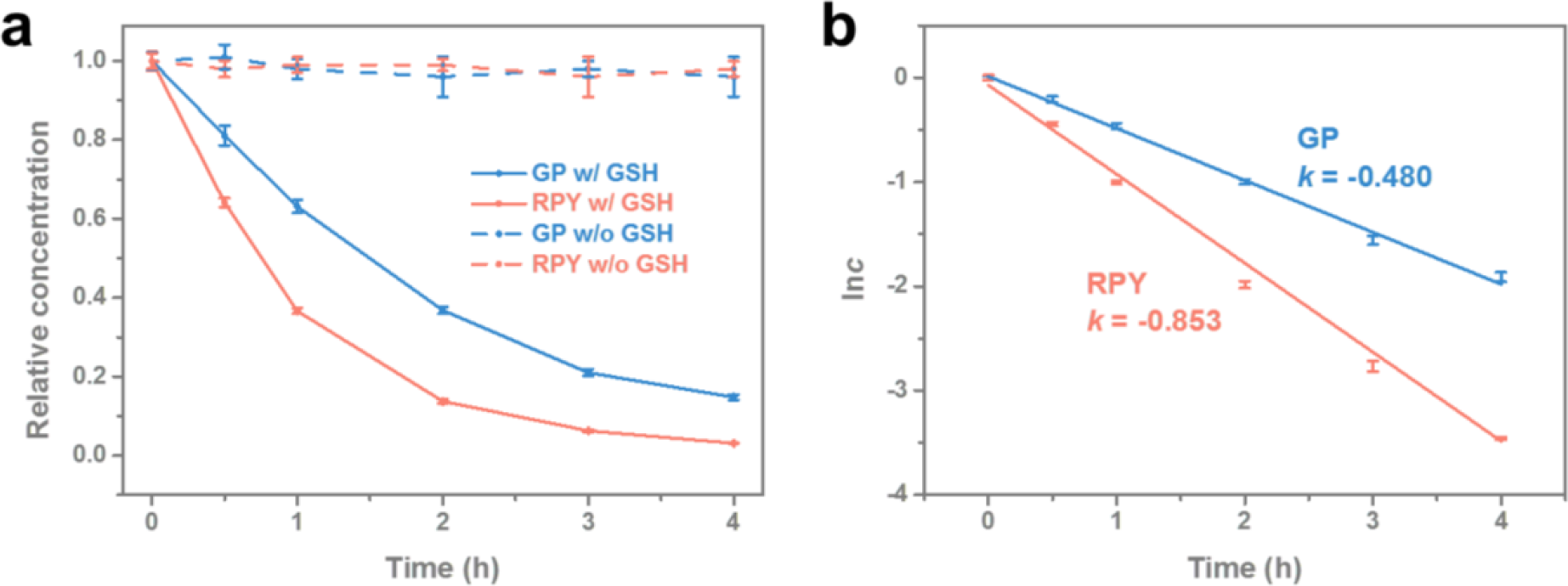
Reduction of the self-immolative moiety of HB*pep*-SP variants induced by GSH measured by HPLC **(a)** Concentration decay of GP and RPY peptide in the presence or absence of 1 mM GSH as a function of time. **(b)** Concentration decay of GP and HPY peptides in natural logarithmic scale in the presence of 1 mM GSH as a function of time. The reaction rate constant *k* was obtained from the slopes of the fitted lines. Data are presented as the mean ± SD of *n* = 3 independent experiments.

**Fig. S7.**
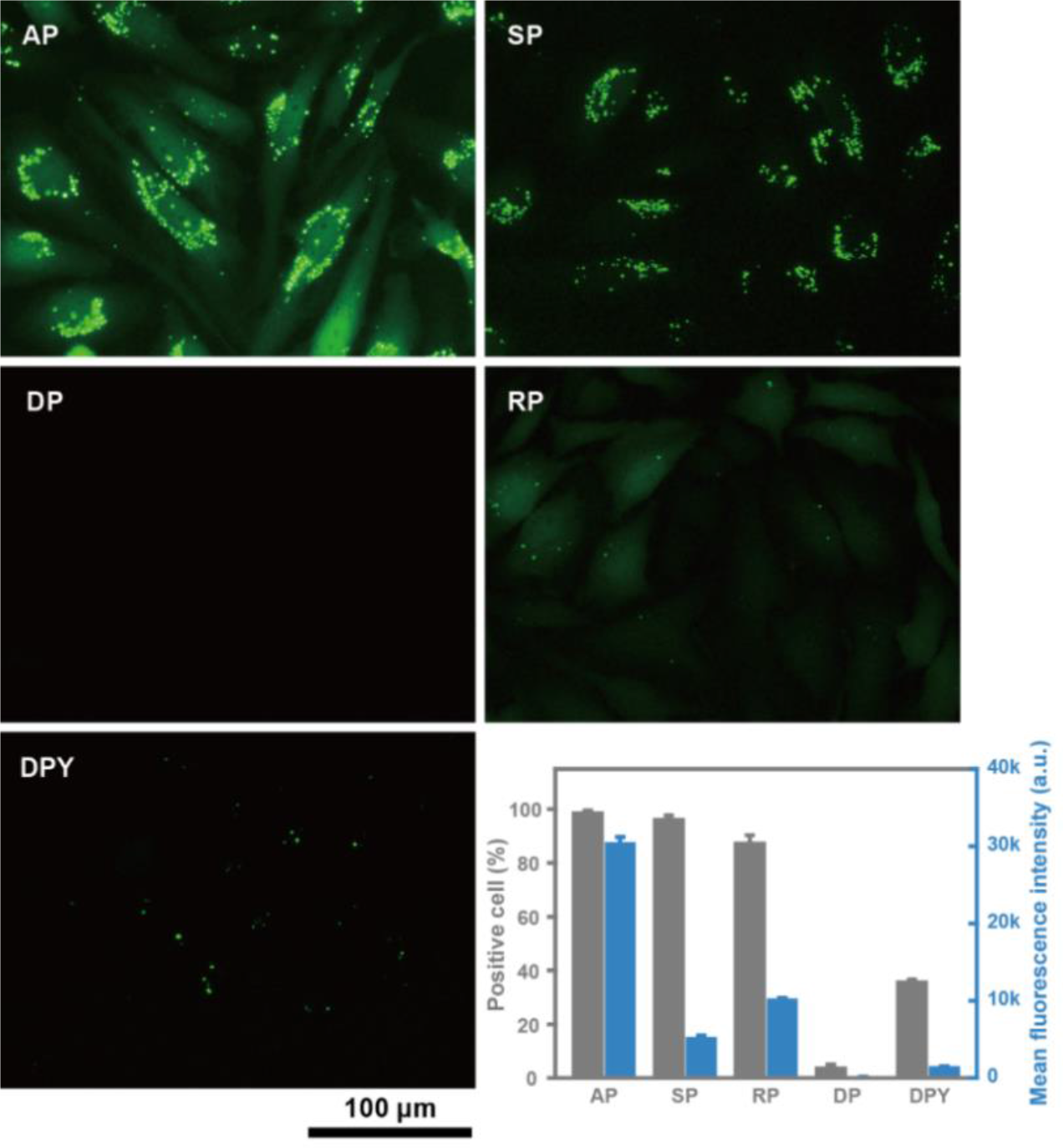
Intracellular delivery of EGFP mediated by HB*pep*-SP CMs variants. Fluorescence micrographs and FACS measurements of HeLa cells treated with EGFP-loaded CMs for 24 hours. Data are presented as the mean ± SD of n = 3 independent experiments.

**Fig. S8.**
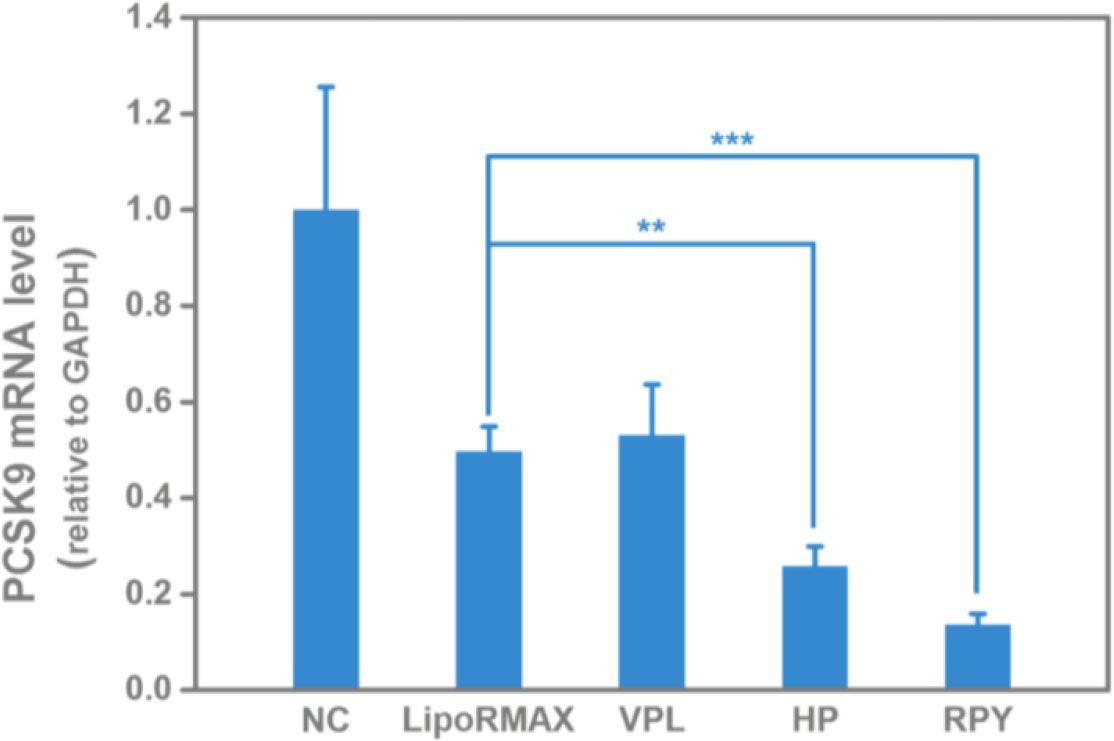
PCSK9 mRNA level in HeLa cells treated with anti-PCSK9 siRNA-loaded CMs variants for 24 hours compared to untreated cells (NC) and LipoRMAX mediated knockdown. Data are presented as the mean ± SD of n = 3 independent experiments; two-sided Student’s t- test, \**P* < 0.05, \*\**P* < 0.01, \*\*\**P*<0.001.

**Fig. S9.**
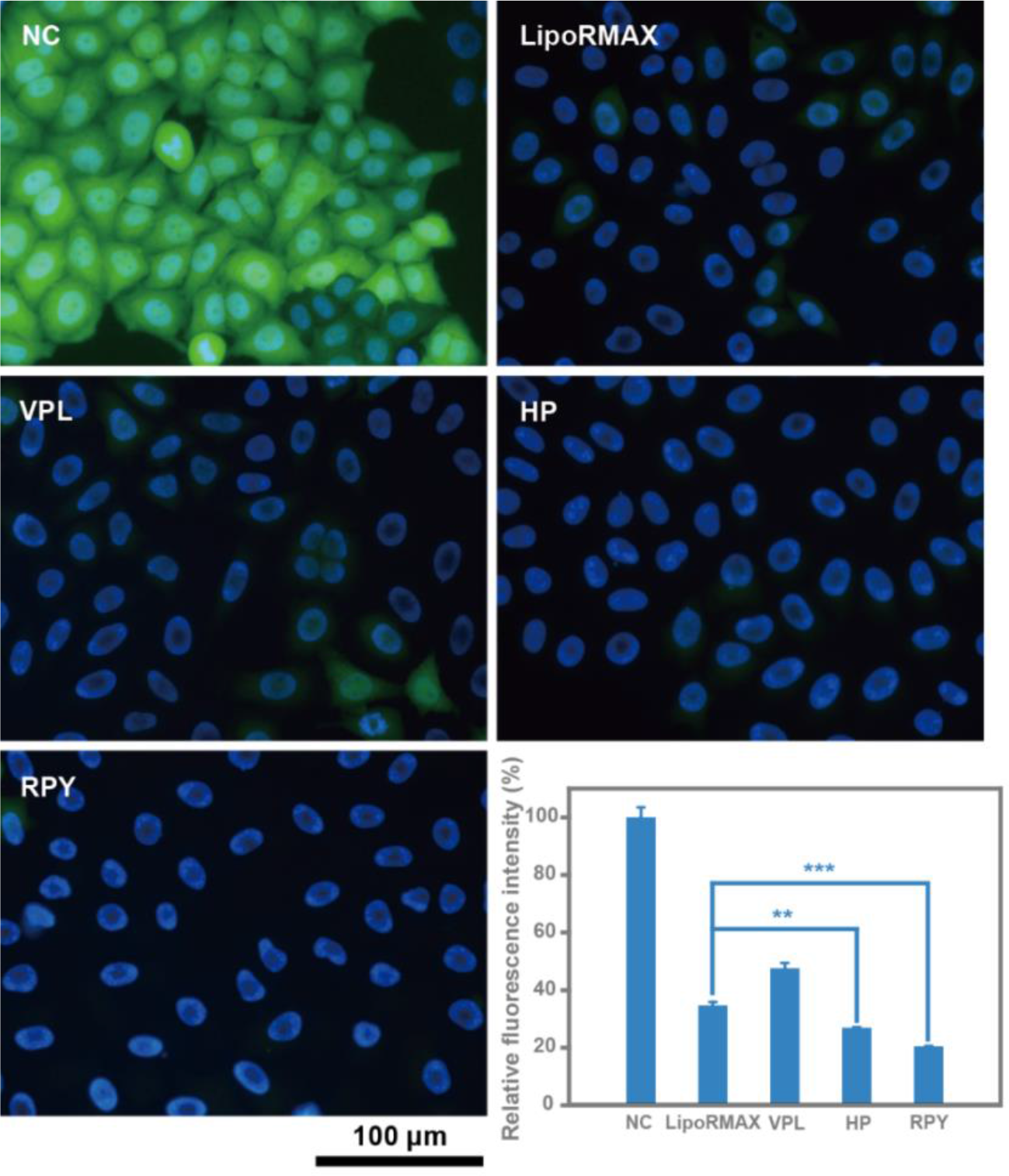
EGFP knockdown mediated by siRNA-loaded CMs variants. Fluorescence micrographs and FACS measurements of HeLa-EGFP cells treated with anti-EGFP siRNA loaded CMs for 24 hours compared to untreated cells (NC) and the commercial reagent LipoRMAX. Data are presented as the mean ± SD of *n* = 3 independent experiments; two-sided Student’s t-test, \**P* < 0.05, \*\**P* < 0.01, \*\*\**P*<0.001.

**Fig. S10.**
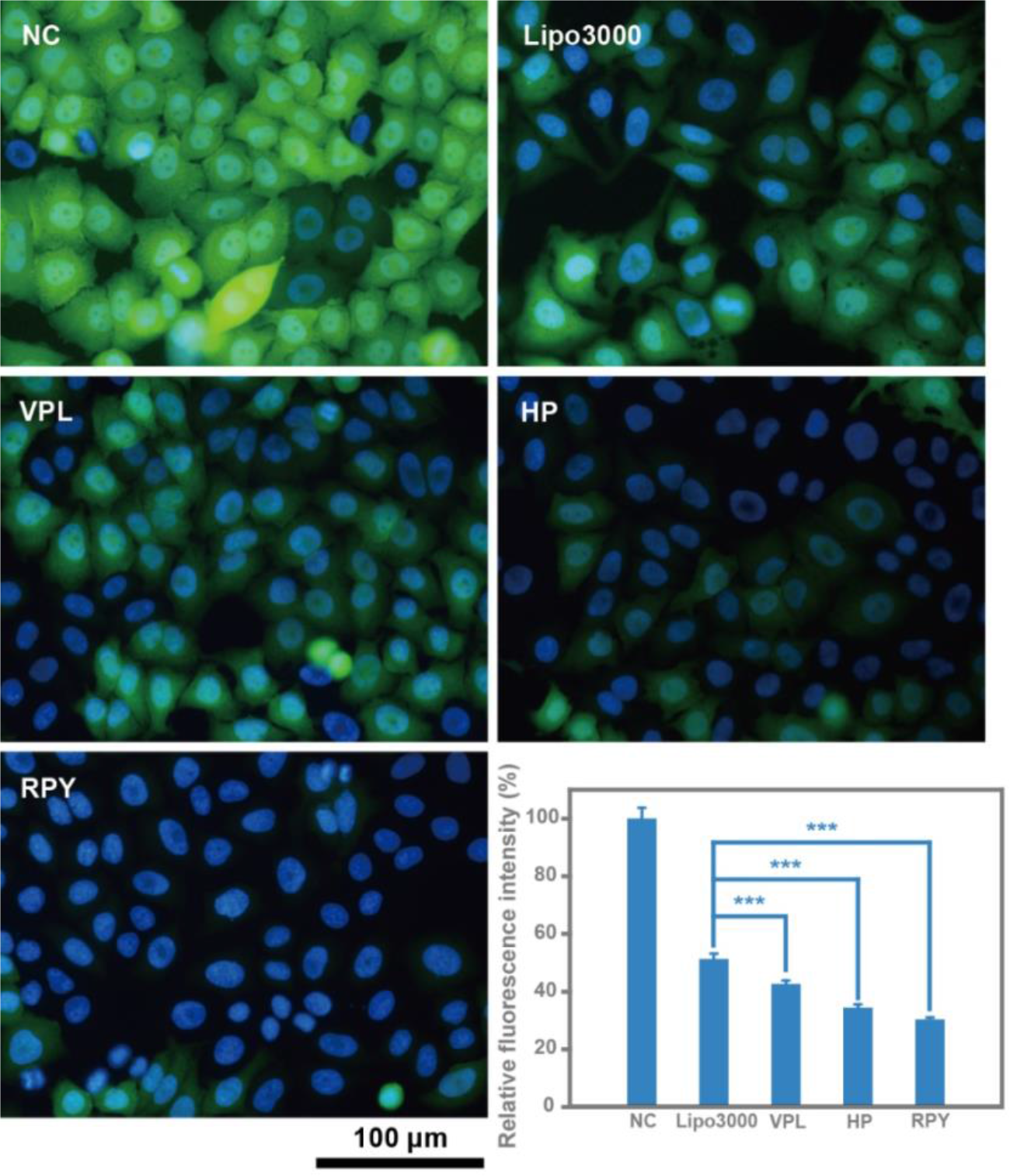
EGFP knockout mediated by CM variants loaded with CRISPR/Cas9 modality. Fluorescence micrographs and FACS measurements of HeLa-EGFP cells treated with CMs loaded with Cas9 mRNA/EGFP-targeting sgRNA for 48 hours compared to untreated cells (NC) and the commercial reagent Lipo3000. Data are presented as the mean ± SD of n = 3 independent experiments; two-sided Student’s t-test, \**P* < 0.05, \*\**P* < 0.01, \*\*\**P*<0.001.

**Fig. S11.**
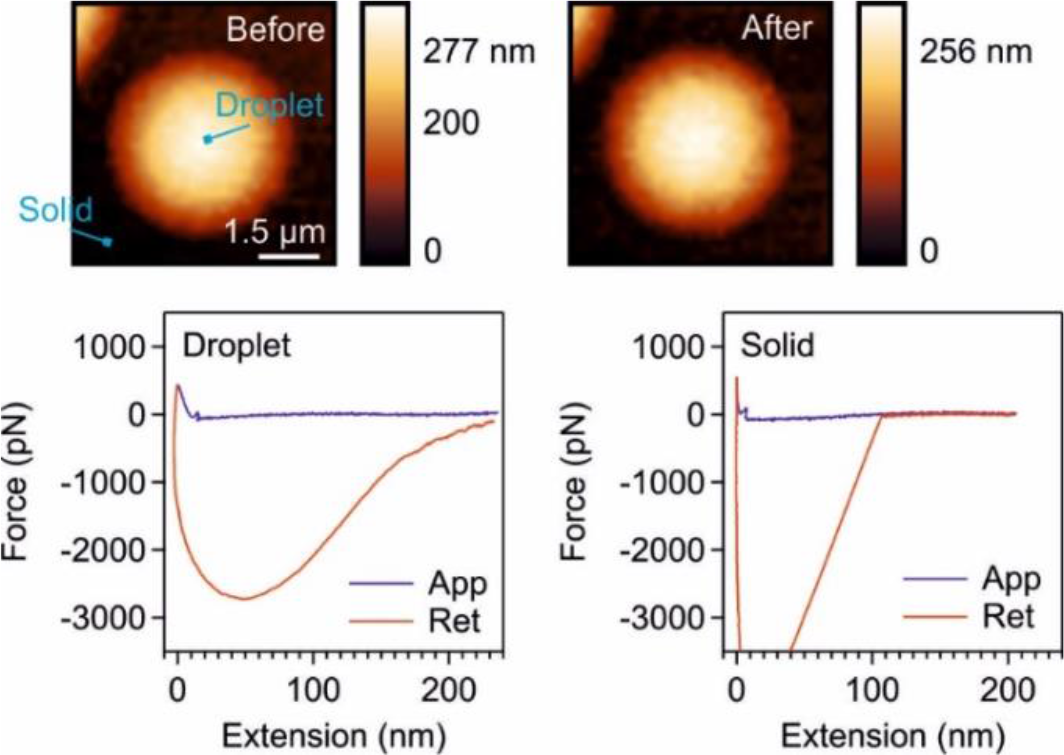
AFM nanoindentation measurements of GP CMs using an AFM tip with stiffness 0.04 N/m. The location of force measurements on CM and solid are indicated.

**Fig. S12.**
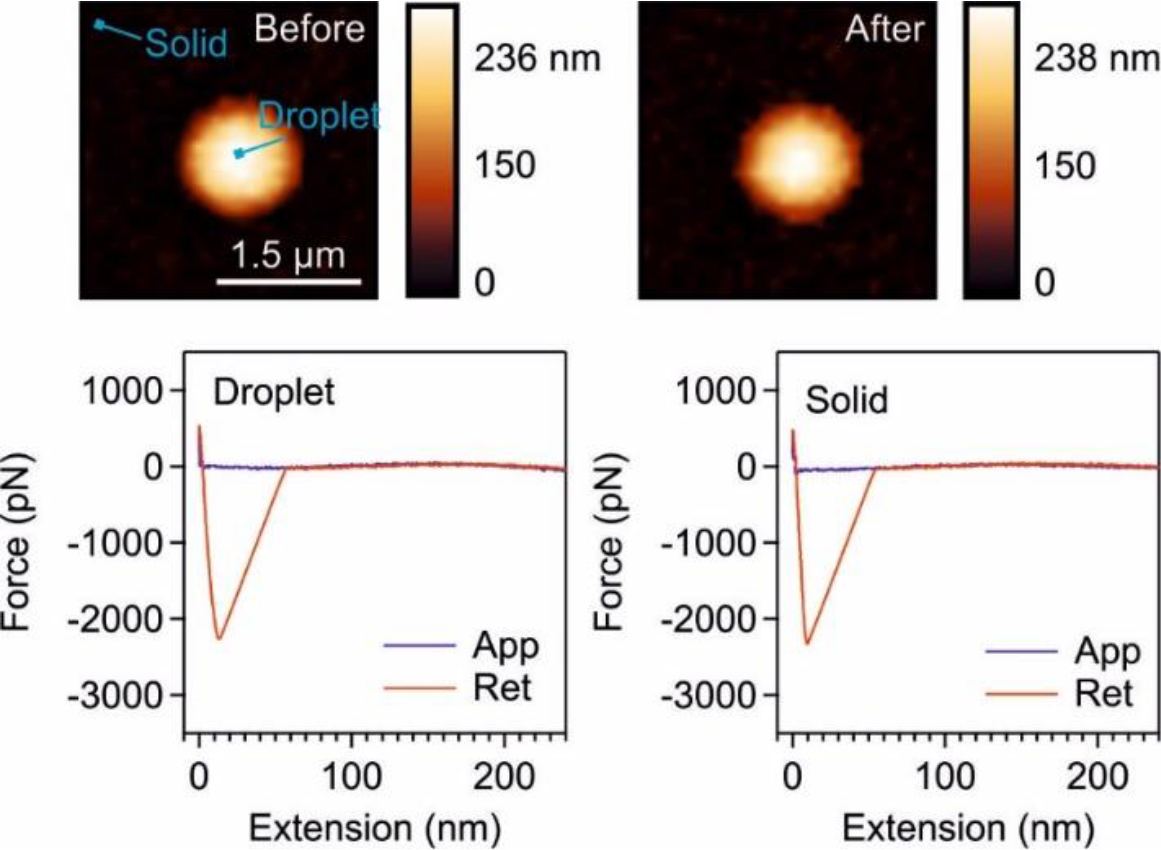
AFM nanoindentation measurement of RPY CMs using an AFM tip with stiffness 0.04 N/m. The location of force measurements on CM and solid are indicated.

**Fig. S13.**
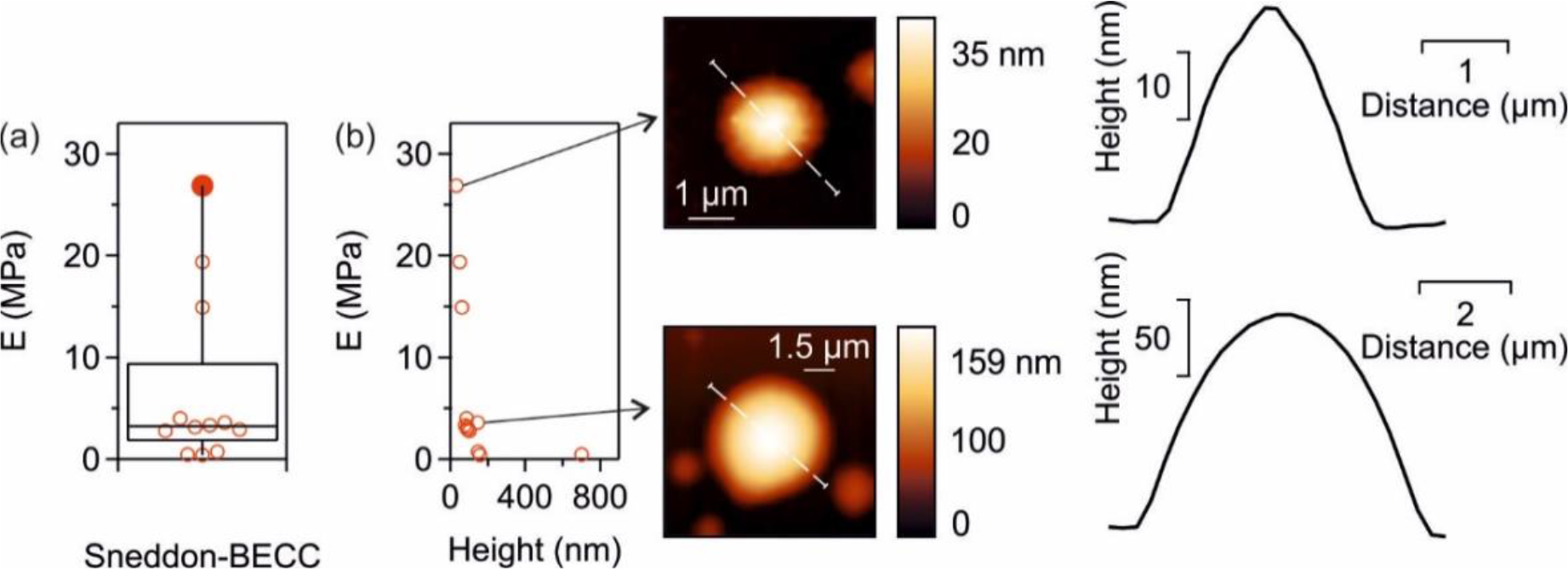
(a) Box plot of the elastic moduli of GP CMs (Igor Pro, WaveMetrics). **(b)** Correlation of elastic moduli with height of GP CMs, together with examples of small- (top) and high- height (bottom) CMs and their respective profiles.

**Fig. S14.**
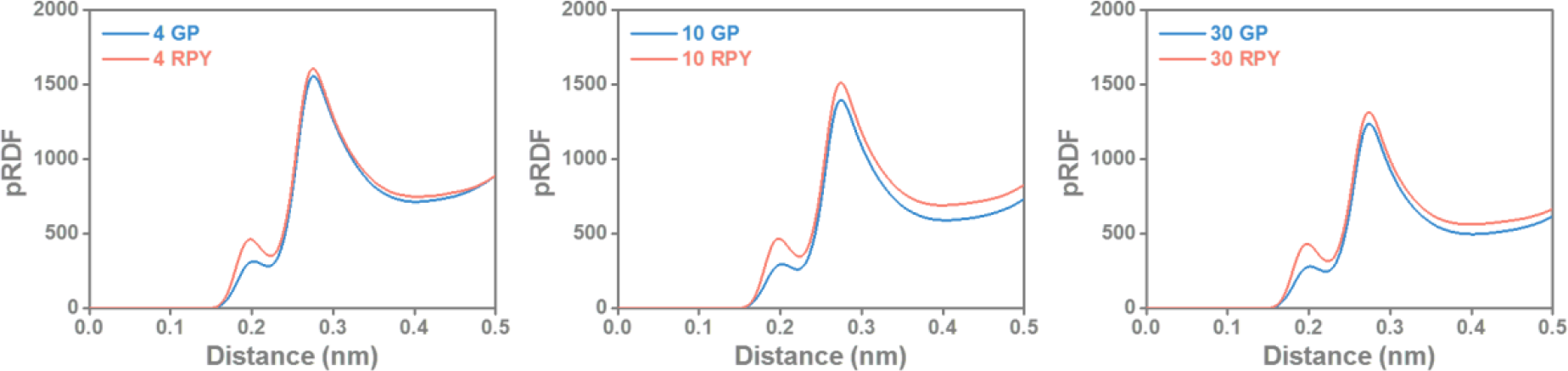
Proximal radial distribution functions of water with respect to the peptides.

**Fig. S15.**
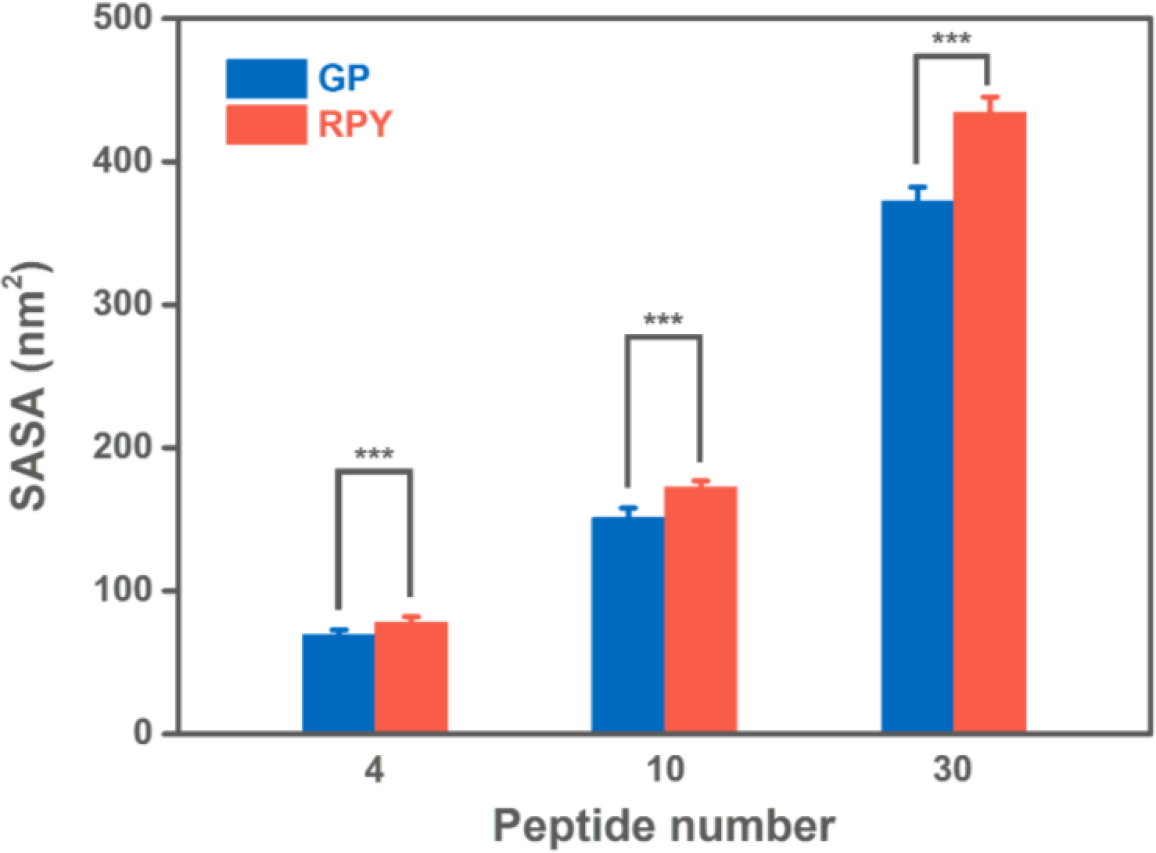
Solvent accessible surface area of clusters with 4, 10 and 30 peptide molecules. SASA was calculated using a probe radius of 0.14 nm. Data are presented as the mean ± SD of *n* = 9000 independent frames; two-sided Student’s t-test, \**P* < 0.05, \*\**P* < 0.01, \*\*\**P*<0.001.

**Fig. S16.**
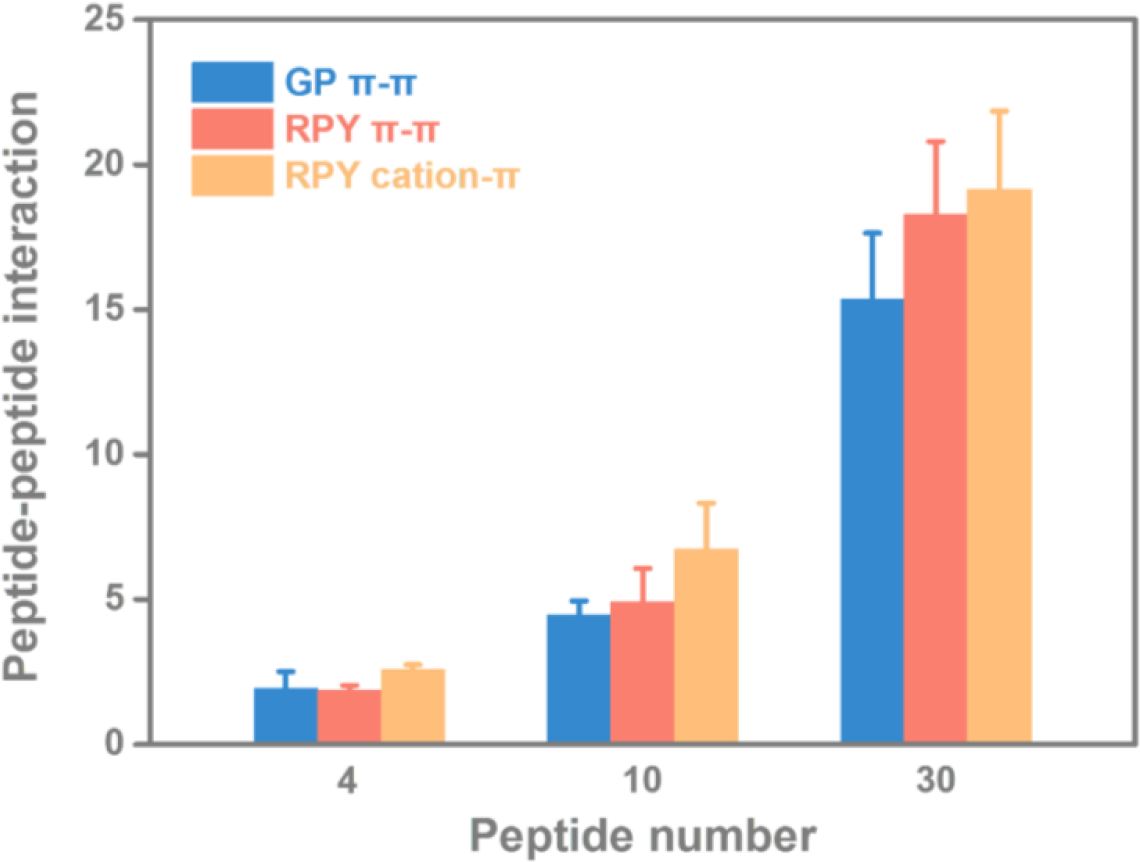
Number of pairs of π-π and cation-π interactions for clusters with different number of peptide molecules. The π-π interaction is defined using a distance cutoff of 0.5 nm between the center of mass of the two aromatic rings and an angle cutoff of 30 degrees between the two aromatic rings. The cation-π interaction is defined using a distance cutoff of 0.5 nm between the center of mass of the guanidinium group of Arg and the center of mass of the aromatic ring, plus an angle cutoff of 30 degrees between the guanidinium plane and the aromatic ring.

**Fig. S17.**
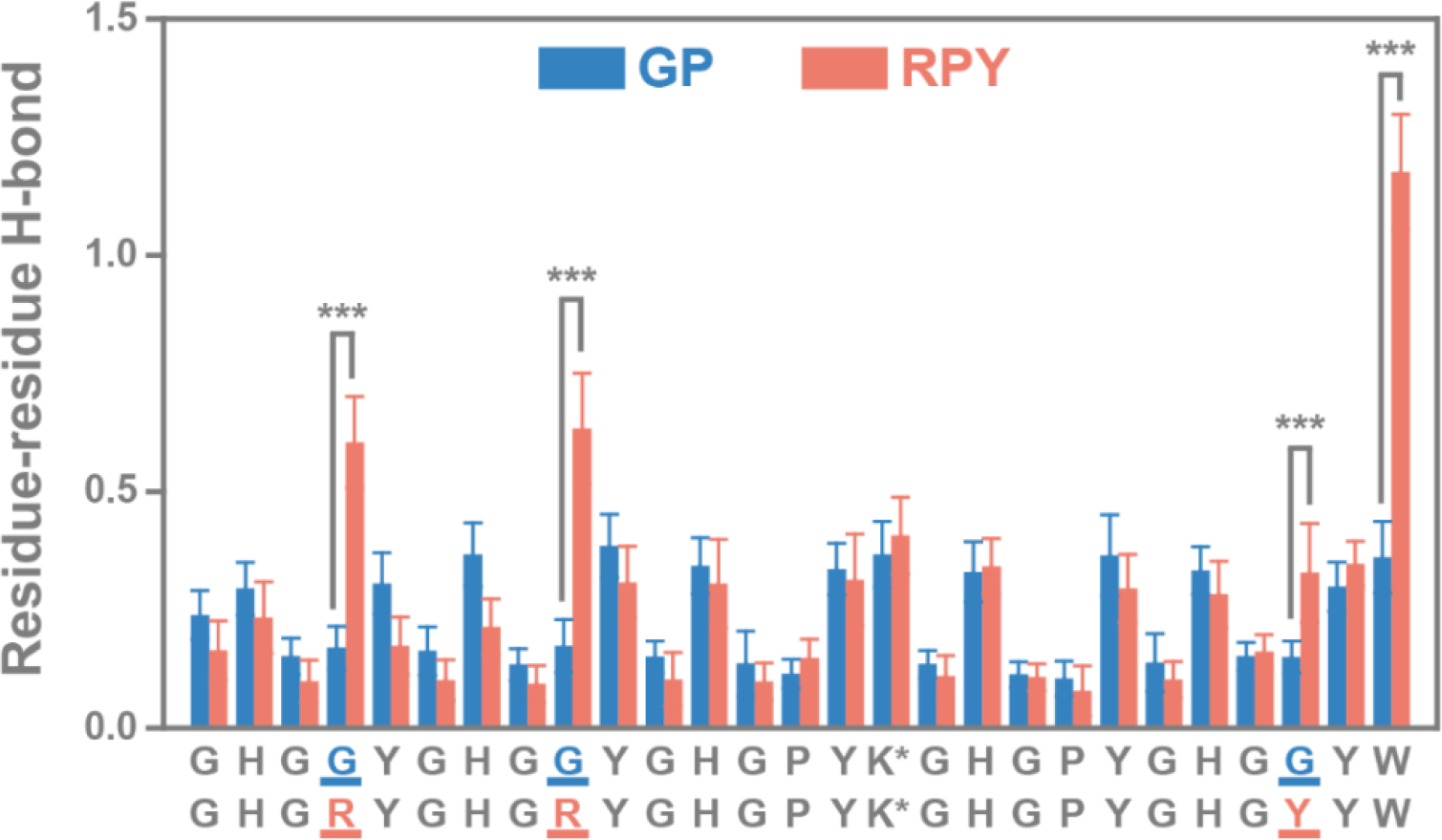
Average number of hydrogen bonds between individual residues in each cluster (30 peptides). Data are presented as the mean ± SD of *n* = 9000 independent frames; two-sided Student’s t-test, \**P* < 0.05, \*\**P* < 0.01, \*\*\**P*<0.001.

**Fig. S18.**
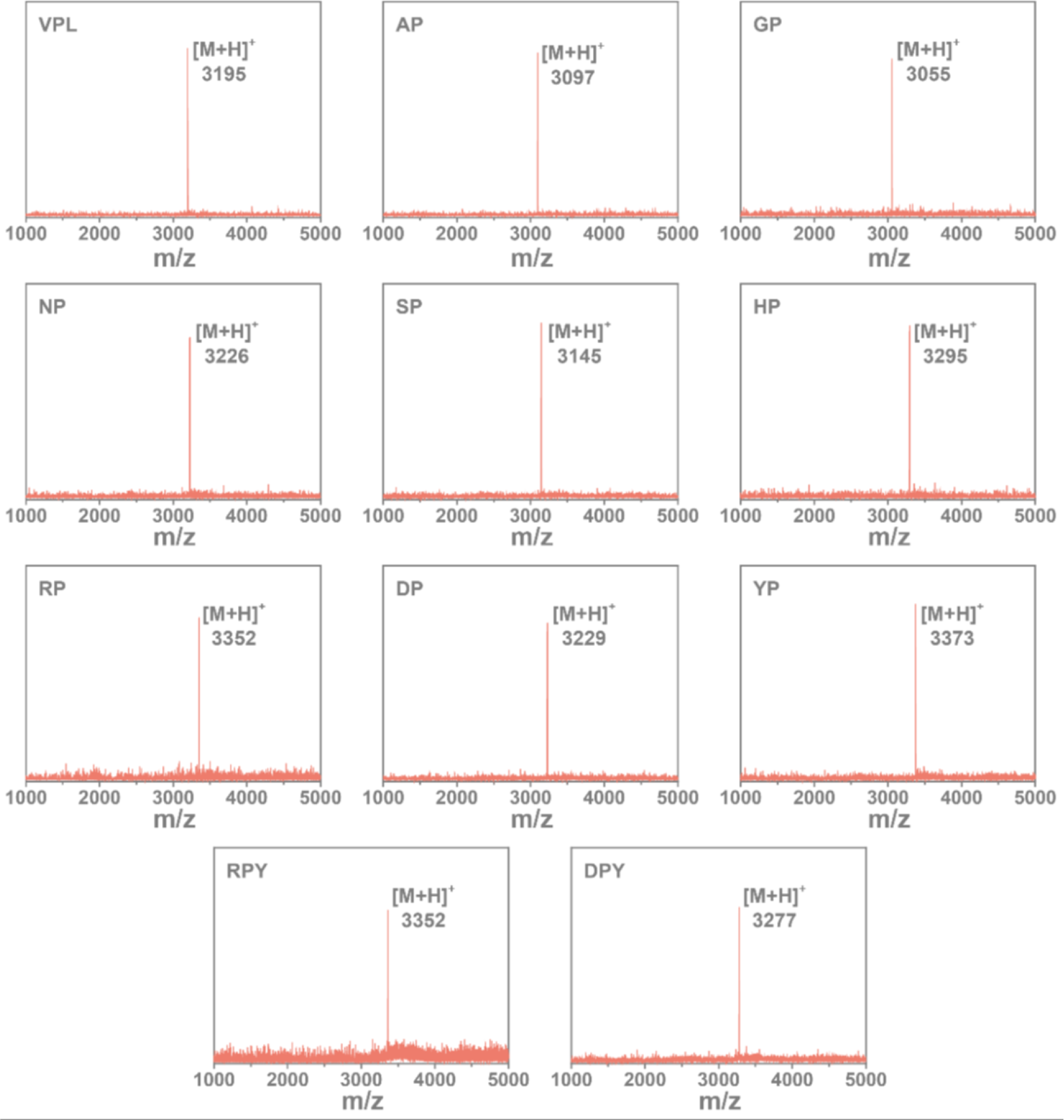
MALDI-TOF spectra of purified HB*pep*-SP variants.

**Fig. S19.**
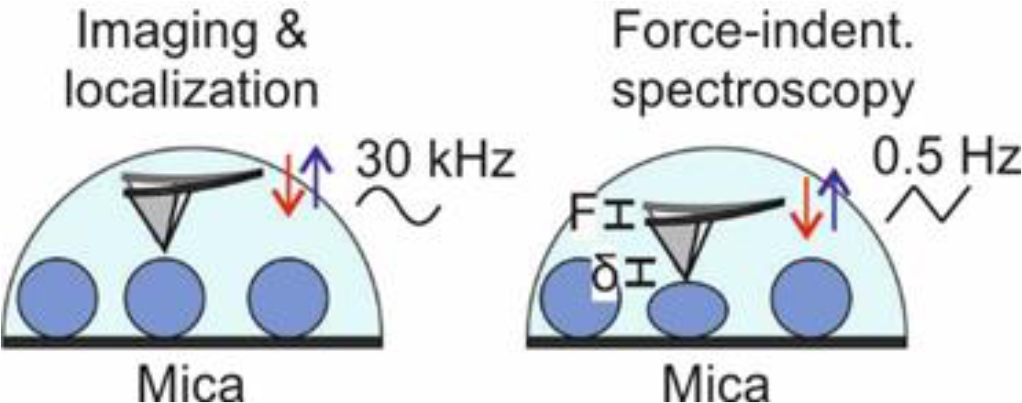
The schematic of AFM measurements on CM variants.

**Fig. S20.**
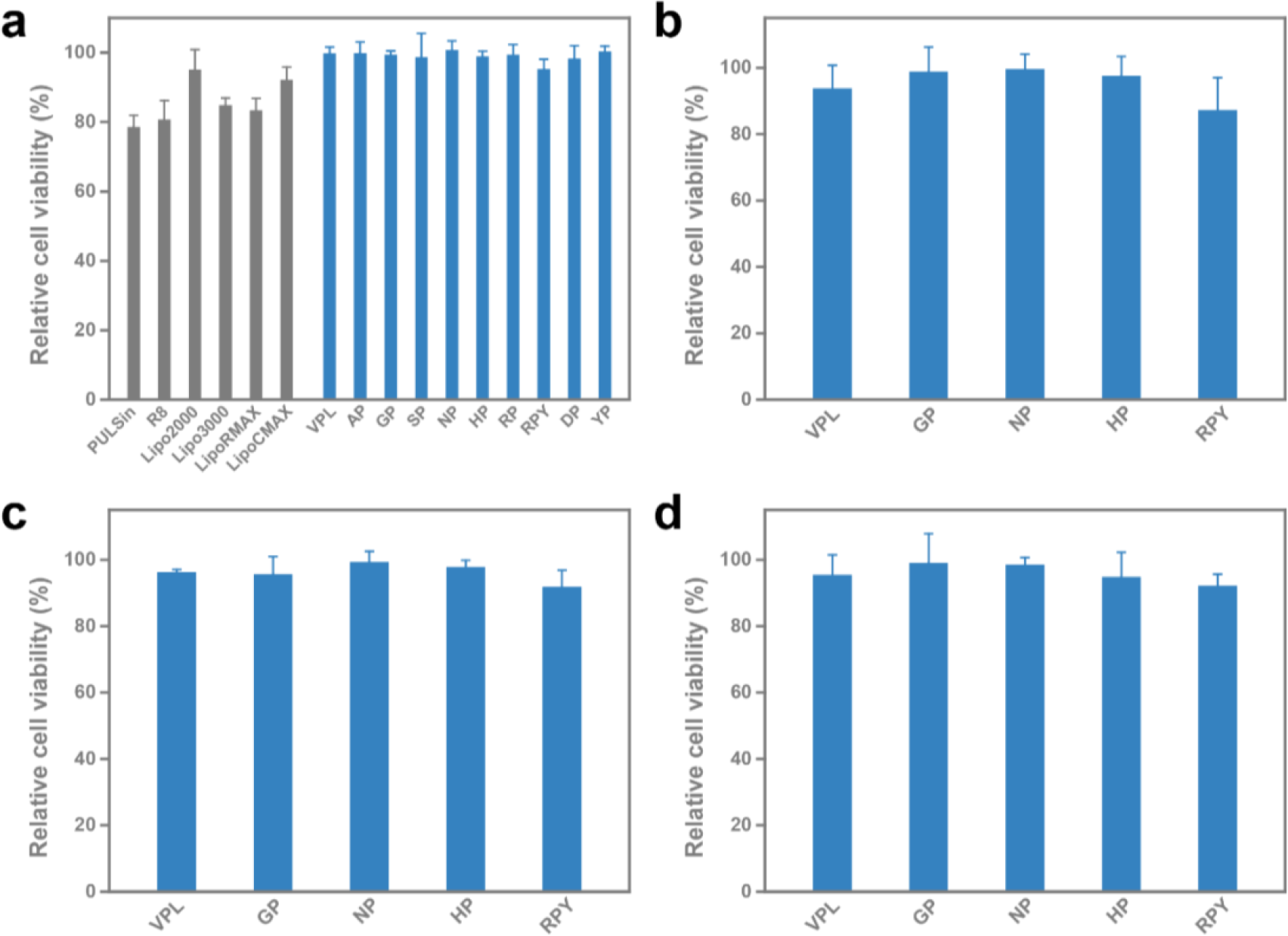
Cytotoxicity of HB*pep*-SP CMs variants. **(a)** Relative cell viability of HeLa cells treated with CMs variants and commercially available transfection reagents. **(b)** Relative cell viability of HFF cells treated with representative CMs variants. **(c)** Relative cell viability of Jurkat T-cells treated with representative CMs variants. **(d)** Relative cell viability of RAW 264.7 cells treated with representative CMs variants. Data are presented as the mean ± SD of n = 3 independent experiments.

**Table S1.**
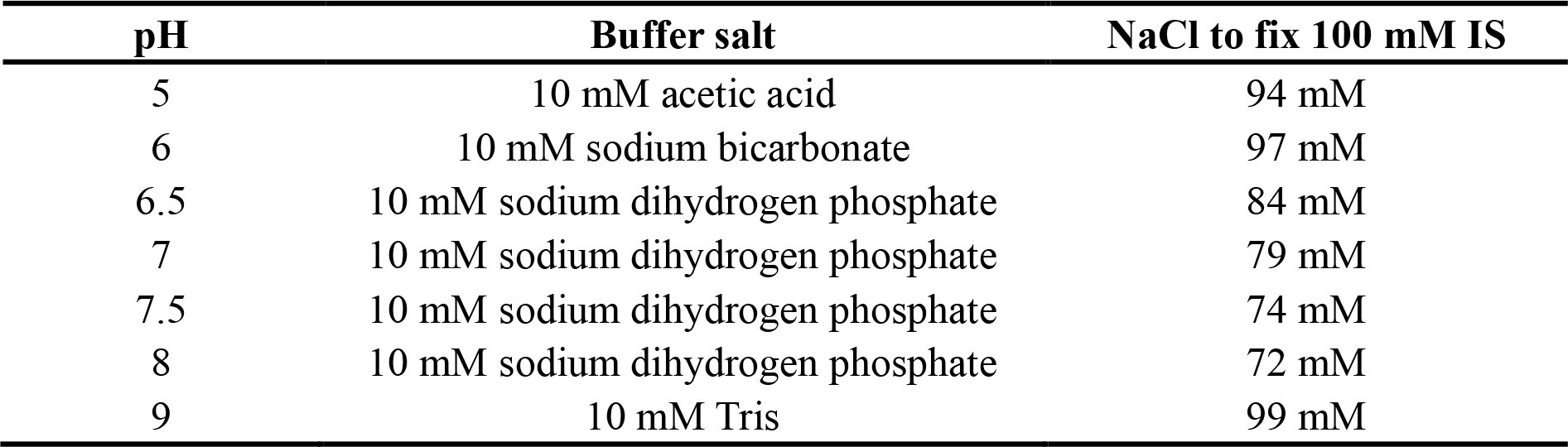
The details of buffers used for LLPS study of HBpep-SP variants.

**Table S2.**
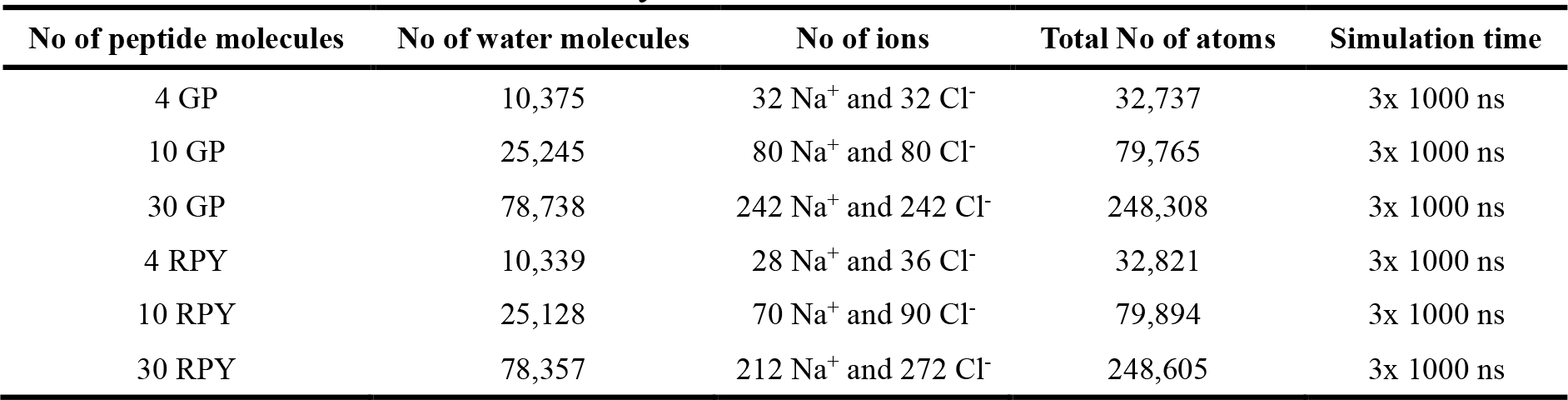
Details of each simulation system.

**Table S3.**
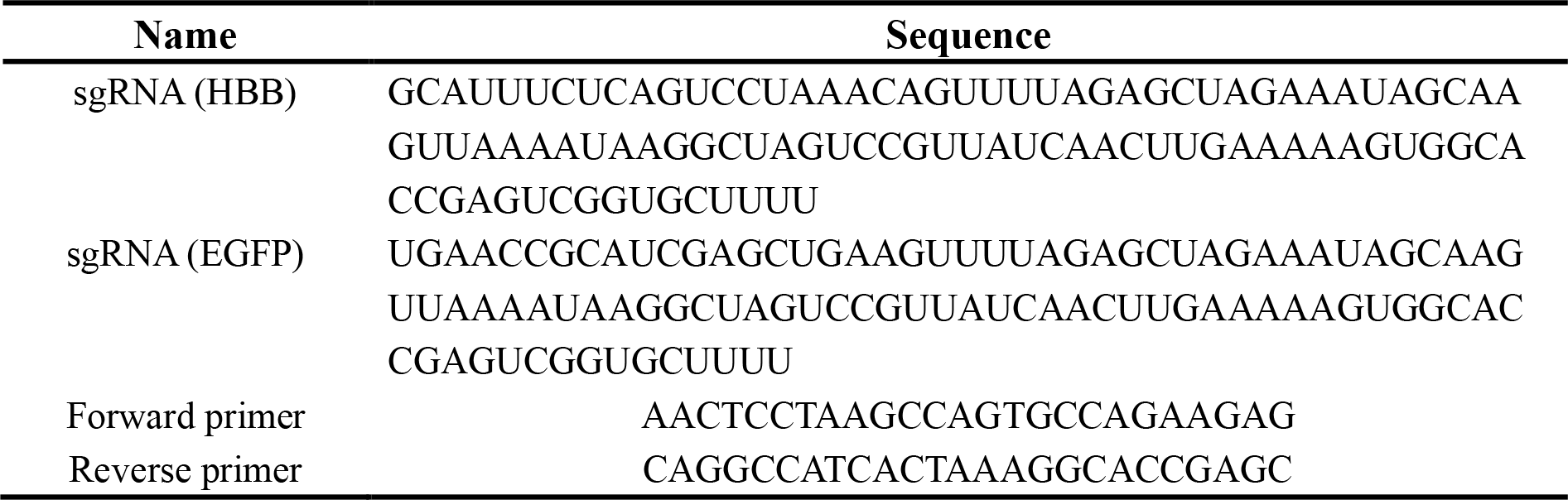
The sequence of sgRNA targeting HBB and EGFP, and primers for amplifying HBB locus.

